# Metabolic and Process Engineering for Microbial Production of Protocatechuate from Xylose with *Corynebacterium glutamicum*

**DOI:** 10.1101/2021.02.12.430943

**Authors:** Mohamed Labib, Jonas Görtz, Christian Brüsseler, Nicolai Kallscheuer, Jochem Gätgens, Andreas Jupke, Jan Marienhagen, Stephan Noack

## Abstract

3,4-Dihydroxybenzoate (protocatechuate, PCA) is a phenolic compound naturally found in edible vegetables and medicinal herbs. PCA is of interest in the chemical industry as a building block for novel polymers and has wide potential for pharmaceutical applications due to its antioxidant, anti-inflammatory, and antiviral properties. In the present study, we designed and constructed a novel *Corynebacterium glutamicum* strain to enable the efficient utilization of d-xylose for microbial production of PCA. The engineered strain showed a maximum PCA titer of 62.1 ± 12.1 mM (9.6 ± 1.9 g L^−1^) from d-xylose as the primary carbon and energy source. The corresponding yield was 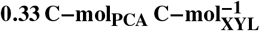, which corresponds to 38 % of the maximum theoretical yield and is 14-fold higher compared to the parental producer strain on d-glucose. By establishing a one-pot bioreactor cultivation process followed by subsequent process optimization, the same maximum titer and a total amount of 16.5 ± 1.1 g was reached. Downstream processing of PCA from this fermentation broth was realized via electrochemically induced crystallization by taking advantage of the pH-dependent properties of PCA. Since PCA turned out to be electrochemically unstable in combination with several anode materials, a three-chamber electrolysis setup was established to crystallize PCA and to avoid direct anode contact. This resulted in a maximum final purity of 95.4 %. In summary, the established PCA production process represents a highly sustainable approach, which will serve as a blueprint for the bio-based production of other hydroxybenzoic acids from alternative sugar feedstocks.

## Introduction

Due to the increasing environmental concerns and the desire for a sustainable bioeconomy there is a growing interest in the production of high-value compounds from renewable feedstocks. Improving the native properties of microorganisms or designing new microbial production hosts is a promising way into that direction. Applications for aromatic compounds, such as the phenolic acid 3,4-dihydroxybenzoate (protocatechuate, PCA), are steadily increasing due to their use in the pharmaceutical and polymer industries.

PCA has a wide range of pharmaceutical applications as an antibacterial, antiviral, antiaging, or antifibrotic agent [1]. Furthermore, its anticancer activity has been reported in terms of an induction of apoptosis of human leukemia cells [2]. Additionally, the antioxidative activity of PCA is based on the counteraction against free radical formation by upregulation of genes encoding enzymes with neutralizing activities [3]. Industrially, the copolymer of PCA and aniline serves as an electrode with high electrochemical potential rendering PCA a precursor of polymers and plastics [4].

PCA naturally occurs as a secondary metabolite in various plant species. For example, it is present in Acaí oil, obtained from the fruit of the Acaí palm (*Euterpe oleracea*) [5]. It is also found as an antifungal agent in the pigmented onion scales of *Allium cepa*, enabling them to resist onion smudge [6]. The flowers of *Hibiscus sabdariffa* contain PCA as an antihypertensive agent [7]. Recently, a selective PCA extraction method for plant material using molecularly imprinted polymers was presented [8]. Its application to the leaves of *Ilex chinensis* Sims yielded 8.46 μg g^−1^. PCA was also purified from the bark of *Terminalia nigrovenulosa* using methanol extraction, followed by fractionation with different solvents [9]. The freeze-dried ethyl acetate fraction resulted in the detection of 1.0 mg mL^−1^ PCA, which is still far too low for an industrial production scale.

Various microorganisms have also shown their potential for PCA biosynthesis. For example, the ubiquitous soil bacterium *Bacillus thuringiensis* excretes PCA into the medium under iron-limiting conditions [10]. Furthermore, *Azotobacter paspali* accumulated PCA upon its cultivation in defined medium containing acetate and d-glucose or sucrose as carbon sources [11]. However, the resulting natural product titers in both cases are very low.

The Gram positive, non-sporulating bacterium *Corynebacterium glutamicum* is widely used in industrial biotechnology for the large scale production of various amino acids such as L-lysine (> 1.4 million t/a) and L-glutamate (> 2 million t/a) [12]. Furthermore, the production of biobased organic acids such as pyruvate, lactate, and succinate has been reported using genetically engineered *C. glutamicum* [13]. The broad spectrum of carbon utilization and plasticity of its metabolism are physical properties that render *C. glutamicum* accessible to manipulation and robust for cultivation under industrial conditions [14].

In a first approach, the L-phenylalanine-producing strain *C. glutamicum* ATCC 21420 was further modified to express the gene *ubiC* coding for chorismate pyruvate lyase from *Escherichia coli*, which enabled the formation of 7.4 mM PCA from d-glucose after 96 h fed-batch cultivation [17]. Recombinant expression of the gene *vanAB* encoding for the heterodimeric vanillate O-demethylase from *Corynebacterium efficiens* NBRC 100395 in the same parental strain enabled the bioconversion of 16.0 mM ferulic acid to 6.91 mM PCA after 12 h of fed-batch cultivation [18]. Noteworthy, *C. glutamicum* is capable of utilizing PCA as sole carbon and energy source [19, 20] and in both approaches followed by Okai and co-workers, the natural PCA catabolism of *C. glutamicum* was not inactivated. Most recently, a high-titer production process for PCA from d-glucose was presented [21]. For reaching the reported titer of 82.7 g L^−1^ PCA a two-step fermentation approach was applied. Cells were first grown to high densities using a complex medium, followed by a manual centrifugation step to harvest the cells for subsequent biotransformation on d-glucose.

Refined d-glucose is the major substrate for glycolytic pathways and is thus preferentially used as a feedstock for biotechnological production using engineered microorganisms [22, 23]. Plants such as sugarcane or sugar beets are an important source of d-glucose and also grow on farmland suitable for food production. Therefore, the use of d-glucose for bioprocesses is considered competitive with human food and may increase commodity prices [24, 25]. By constrast, the C5 sugar d-xylose is the second most abundant fraction of lignocellulosic biomass generated as waste from agricultural, pulp and paper industry [26, 27]. Consequently, it is a costeffective renewable carbon source compared to typically used hexoses. To date, all existing bioprocesses for PCA production use d-glucose or mixtures containing different hexoses. High-yield microbial production of PCA from renewable feedstocks requires a proper protocol for product extraction from the cultivation broth, in particular when considering that the microbial producer forms compounds with a similar polarity as part of its central metabolism. For example, ultrafiltration [28], adsorption [29], are energy- and time-consuming options for PCA recovery from fermentation medium [30]. Recently, tri-*n*-butyl phosphate was used as a reactive extraction agent to form a complex with protonated PCA in the aqueous phase that can be extracted using an organic phase [31–34]. Moreover, an *in situ* product removal concept using reactive extraction was suggested to prevent product inhibition during the fermentation process [33]. Nevertheless, this concept is limited for fermentation processes with a pH below the respective *pK*_*a*,*COOH*_ value of 4.48 [35]. Additionally, the solubility of organic solvents in the aqueous phase can negatively affect the fermentation process [36]. Most recently, a promising downstream processing strategy for succinic acid was developed that avoided waste salt production and the use of an organic phase by inducing the pH shift electrochemically and crystallization of the target component [37]. This purification protocol can be adopted for the separation of PCA from fermentation medium.

In this study, we established a one-pot production process for bio-based PCA that enables efficient decoupling of biomass and product formation utilizing d-glucose and d-xylose as complementary carbon substrates. By combining *in silico* strain design with targeted metabolic engineering and process development including downstream processing, a sustainable and industrially relevant production process for PCA could be developed.

## Results and discussion

### *In silico*-guided improvement of PCA-producing *C. glutamicum* strains

The most urgent prerequisite for exploiting *C. glutamicum* as producer of PCA is the abolishment of a catabolic pathway for PCA present in the wild-type strain. A respective platform strain, in which most of the peripheral and central degradation pathways for aromatic components have been abolished, was constructed recently [16]. Additional engineering work focused on improved d-glucose import through deregulation of the gene for the glucose/*myo*-inositol permease IolT1. Further modifications were introduced to improve the carbon flux into the shikimate pathway. These included a reduced flux towards the tricarboxylic acid (TCA) cycle by reduction of native citrate synthase activity and expression of a codon-optimized version of the gene *aroF*\* from *E. coli*. This gene codes for an L-tyrosine feedback-resistant 3-deoxy-d-arabinoheptulosonate-7-phosphate (DAHP) synthase catalyzing the initial and committed step in the shikimate pathway. In combination with overexpression of native genes coding for 3-dehydroshikimate dehydratase QsuB and transketolase Tkt, the constructed strain *C. glutamicum* De-lAro5 C7 P_O6_-*iolT1* pMKEx2_*aroF*\*_*qsuB* pEKEx3_*tkt* (abbreviated as PCA_GLC_) produced 13 mM PCA in shake flask cultures starting from 4 % (w v^−1^) d-glucose as sole carbon and energy source in defined CGXII medium [16].

In our study, flux balance analyses were carried out to identify optimal PCA production routes and to derive further promising genetic engineering targets ultimately leading to improved PCA production in *C. glutamicum*. Starting from d-glucose as sole carbon and energy source a maximum PCA yield of 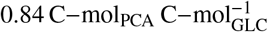 was predicted, which could theoretically be realized under aerobic, growth-decoupled conditions (Figure 1A). The inactivation of pyruvate kinase is predicted as target to avoid loss of the PCA precursor PEP, which is mostly converted to pyruvate and subsequently to the TCA cycle-fueling substrate acetyl-CoA. In fact, deletion of the corresponding *pyk* gene has already been tested for the production of 4-hydroxybenzoate from d-glucose with *C. glutamicum* [38]. In this case, however, the resulting increase in product yield was only 1 %. This is likely due to the fact that the active (i.e. non-abolished) PTS-dependent uptake of d-glucose results in the formation of one mole of pyruvate per mol of imported d-glucose, which would have to be recycled to recover the equivalent amount of PEP for PCA synthesis. In *C. glutamicum*, this could theoretically be achieved by a reaction sequence involving pyruvate carboxylase and phosphoenolpyruvate carboxykinase (PEPCK) (cf. Figure 1A). Although PEPCK is reported to be also active under glycolytic conditions [39], its high net reverse operation under *in vivo* conditions is thermodynamically unfavorable.

**Fig. 1.**
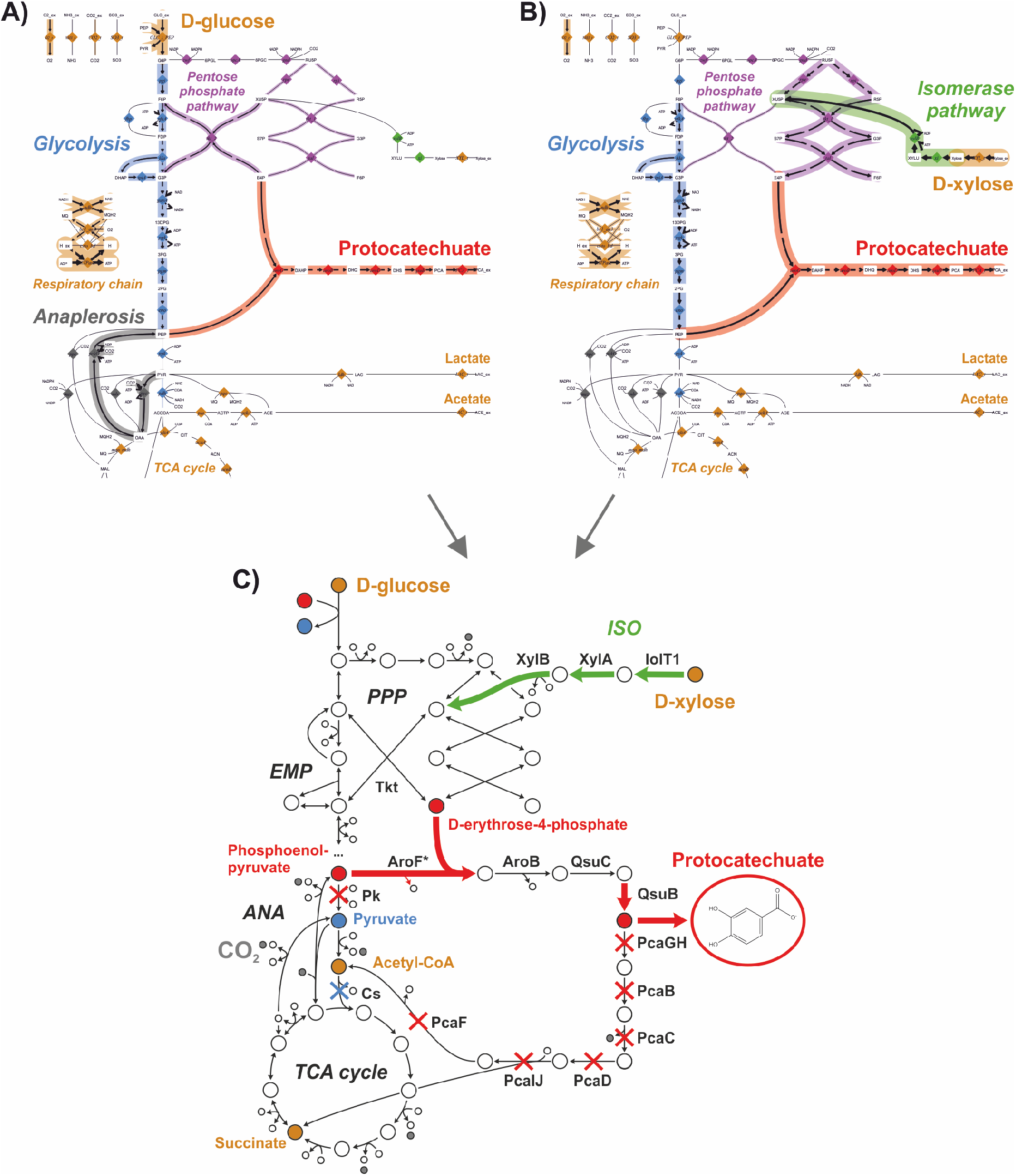
*In silico* strain design and resulting metabolic engineering of *C. glutamicum* for PCA production. A) Optimal PCA production route from d-glucose following PTS-coupled uptake. B) Optimal PCA production route from d-xylose by utilizing the isomerase pathway. In both cases, the colored reactions represent active steps and the thickness of each arrow corresponds to its flux value in relation to the uptake rate. C) Selected engineering targets for *C. glutamicum* enabling production of PCA from d-glucose and d-xylose in a one-pot process. Highlighted arrows represent reactions steps that are enforced through plasmid-based (over)expression or targeted inactivation of native gene regulation. Red crosses represent gene deletions while the blue cross represents a targeted down-regulation of gene expression (to 10 % residual activity compared to the wild-type strain). EMP: EmbdenMeyerhof-Parnas pathway (most common glycolytic pathway), PPP: pentose phosphate pathway, TCA: tricarboxylic acid cycle (citrate cycle), Tkt: transketolase, Pk: pyruvate kinase, Cs: citrate synthase.

Alternatively, a non-PTS route for d-glucose phosphorylation via ATP-dependent hexokinase / glucokinase is conceivable, which would circumvent the loss of PEP and even enable the maximum theoretical yield of 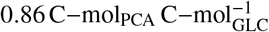. Kogure et al. demonstrated high-yield production of shikimate with a PTS-deficient *C. glutamicum* strain and combined expression of *ppgK* (encoding polyphosphate glucokinase) and *iolT1* (encoding the permase IolT1) [40]. The authors also tested the additional deletion of the *pyk* gene to minimize loss of PEP, but the resulting strain was impaired in growth. To circumvent this problem, a tunable downregulation of the pyruvate kinase activity was proposed that should allow a first growth phase on d-glucose, followed by growth-decoupled production phase [40]. However, such an inducible mechanism, which inhibits the activity of already synthesized proteins is not yet known.

Noteworthy, when using d-glucose as a carbon source, the combined operation of the oxidative and reductive part of the pentose phosphate pathway is not the preferred route for supply of erythrose-4-phosphate (E4P, the second substrate of DAHP synthase besides PEP), since this would result in a loss of carbon in form of CO_2_. Instead, operation of the transketolase 2 enzyme in the direction of E4P formation is required (cf. Figure 1A), which could be supported by additional overexpression of the *tkt_1* gene (cf. Table 1).

**Table 1.**
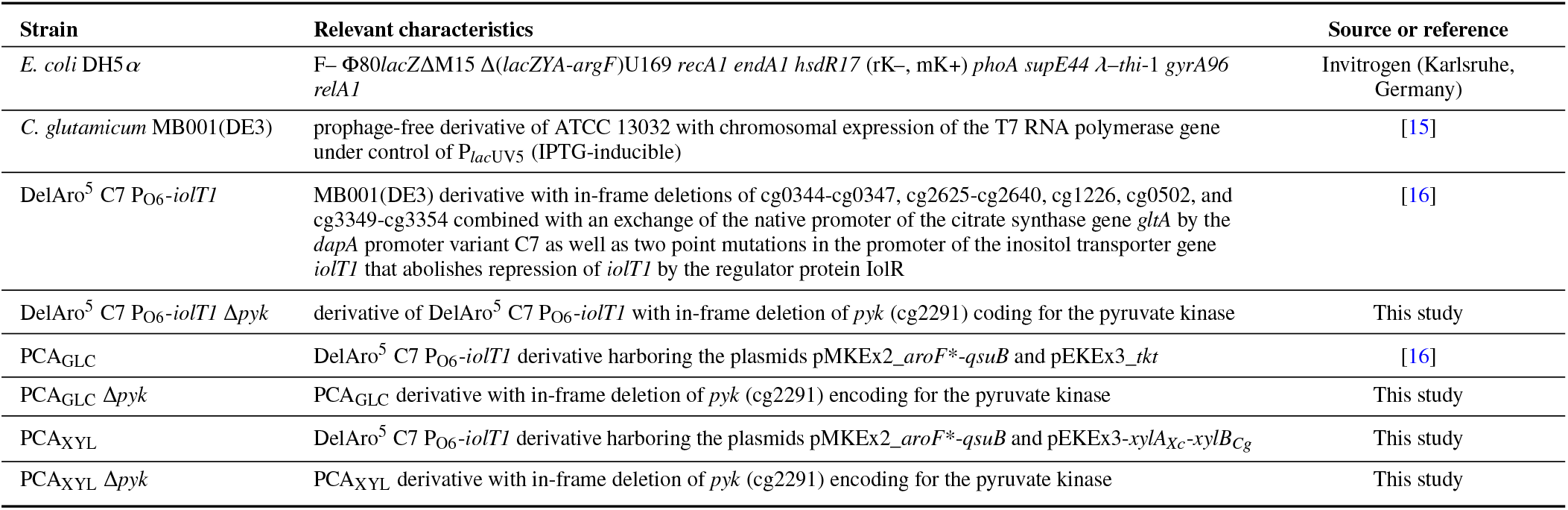
Strains used in this study.

Taking all these aspects into consideration, we decided to follow a different growth-decoupled production strategy for PCA that relies on the utilization of one primary carbon substrate (i.e. d-glucose) for biomass production and another one (i.e. d-xylose) for product formation. By introducing the bacterial isomerase pathway into the native metabolic network of *C. glutamicum*, the non-PTS substrate d-xylose could be converted to PCA with the same maximum theoretical yield of 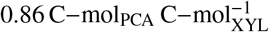 (Figure 1B). While growth-decoupling and PEP accumulation can be realized through *pyk* deletion (without interference of growth on d-glucose), the increased supply of E4P is also guaranteed by using the isomerase pathway for utilization of d-xylose.

Following the predictions of our initial *in silico* study aiming to avoid flux of PEP towards the TCA cycle, we deleted the gene coding for the pyruvate kinase (*pyk*) in our parental producer strain PCA_GLC_ yielding *C. glutamicum* DelAro5 C7 P_O6_-*iolT1* Δ*pyk* pMKEx2_*aroF*\*_*qsuB* pEKEx3_*tkt* (referred to as PCA_GLC_ Δ*pyk*). For establishing the d-xylose-based and growth-decoupled production of PCA, we implemented the isomerase pathway for the degradation of d-xylose in both strains by expressing the heterologous genes coding for xylose isomerase (*xylA*) from *Xanthomonas campestris* and overexpression of the endogenous xylose kinase gene (*xylB*) instead of expressing the transketolase gene (*tkt*). The resulting strain *C. glutamicum* DelAro5 C7 P_O6_-*iolT1* pMKEx2_*aroF*\*_*qsuB* pEKEx3-*xylA_Xc_-*xylB*_Cg_* and its derivative with the additional *pyk* deletion are abbreviated in the following as PCA_XYL_ and PCA_XYL_ Δ*pyk* respectively (cf. Figure 1C).

### Comparative phenotyping of engineered PCA producers

In order to study the impact of d-xylose assimilation and *pyk* deletion on PCA production, all four strains were cultivated in shake flasks in defined CGXII medium, supplemented with either 222 mM d-glucose or 266 mM d-xylose as sole carbon and energy source. Induction of episomal gene expression was achieved by supplementation of the inducer IPTG directly after inoculation of the cultures. All four strains take up d-glucose by the native PTS-system for glucose as well as by the transporter IolT1. The gene coding for the latter is normally not expressed under the chosen cultivation conditions, however, an engineered constitutive expression of the *iolT1* gene in all strains described in this study is ensured by modification of the respective operator/promotor sequence (indicated by P_O6_-*iolT1* in the strain designations). The transporter IolT1 is also responsible for non-PTS uptake of d-xylose [41].

Cultivation of strains PCA_GLC_ and PCA_GLC_ Δ*pyk* on d-glucose showed comparable growth rates of 0.21 h^−1^ and 0.20 h^−1^, respectively, as well as comparable substrate uptake rates of 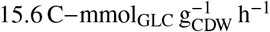 and 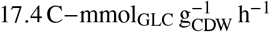 respectively (Figure 2 and Table 2). the PCA production rate 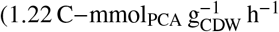, the final PCA titer (7.8 ± 1.6 mM) and PCA yield 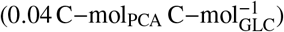 of strain PCA_GLC_ Δ*pyk* were significantly increased compared to its predecessor strain PCA_GLC_. Nevertheless, the PCA yield is still more than one order of magnitude lower than the maximum theoretical yield mentioned above. The strain PCA_XYL_ showed a significantly reduced growth rate (0.11 h^−1^) and hence also final biomass concentration. While specific growth rate (cf. Table 2). The observed slow, but steady growth of strain PCA_XYL_ Δ*pyk* up to an OD_600_ 25 during shake flask cultivation can only be explained by an alternative flux mode for pyruvate supply involving the operation of malic enzyme. In fact, a similar role of malic the l-xylose uptake rate 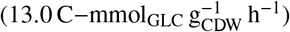 was slightly lower, the overall PCA production performance was further improved in comparison to the PCA_GLC_ *Δpyk* strain. Remarkably, strain PCA_XYL_ *Δpyk* showed the lowest growth rate (0.04 h^−1^), the lowest substrate uptake rate 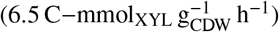 but the highest PCA production rate 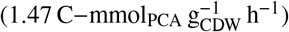 highest final PCA titer (62.1 ± 12.1 mM) and the highest PCA yield 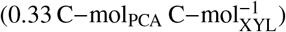of all four strains. Overall, strain PCA_XYL_ Δ*pyk* demonstrated a strong increase in the production-related performance indicators in comparison to the previously published PCA producer strain PCA_GLC_ [16]. In particular, the PCA yield of strain PCA_XYL_ Δ*pyk* corresponds to 38 % of the maximum theoretical yield, which is already acceptable from an industrial perspective for a *de novo* produced benzoic acid.

**Table 2.** Performance indicators of engineered *C. glutamicum* PCA producer strains obtained from batch and fed-batch cultivation experiments. Maximum rate estimation was performed by fitting suitable bioprocess models to the time-dependent measurements covering the productive metabolic states. Mean values and 90 % confidence bounds are derived directly from replicate measurements or via the model-based approach following parametric bootstrapping (see Supplementary Information for more details).

| Strain/ condition | Batch shake flask (*n* = 3) |  |  |  | Fed-batch bioreactor (*n* = 2) |  |
| --- | --- | --- | --- | --- | --- | --- |
| | PCA<sub>GLC</sub> | PCA<sub>GLC</sub> $\Delta pyk$ | PCA<sub>XYL</sub> | PCA<sub>XYL</sub> $\Delta pyk$ | PCA<sub>XYL</sub> $\Delta pyk$<br>cond A | cond B<sup>a</sup> |
| Final OD<sub>600</sub> [-] | 74.2 ± 8.5 | 67.7 ± 6.9 | 41.0 ± 4.0 | 24.7 ± 6.5 | 11.6 ± 7.5 | 50.8 ± 9.5 |
| Final PCA titer [mM] | 3.5 ± 1.2 | 7.8 ± 1.6 | 12.0 ± 1.1 | 62.1 ± 12.1 | 18.0 ± 12.4 | 61.7 ± 4.0 |
| Maximum growth rate [h<sup>-1</sup>] | 0.21 [0.205, 0.208] | 0.20 [0.198, 0.202] | 0.11 [0.105, 0.108] | 0.04 [0.035, 0.042] | 0.40 [0.404, 0.422] | - |
| Maximum D-glucose uptake rate [C–mmol<sub>GLC</sub> g<sub>CDW</sub><sup>-1</sup> h<sup>-1</sup>] | 15.6 [15.57, 16.00] | 17.4 [17.38, 17.82] | - | - | 19.4 [19.47, 21.97] | - |
| Maximum D-xylose uptake rate [C–mmol<sub>XYL</sub> g<sub>CDW</sub><sup>-1</sup> h<sup>-1</sup>] | - | - | 13.0 [12.84, 13.61] | 6.5 [5.95, 7.17] | 3.5 [3.43, 3.79] | - |
| Maximum PCA production rate [C–mmol<sub>PCA</sub> g<sub>CDW</sub><sup>-1</sup> h<sup>-1</sup>] | 0.78 [0.765, 0.832] | 1.22 [1.210, 1.279] | 1.43 [1.257, 1.776] | 1.47 [1.388, 1.625] | 0.53 [0.315, 0.793] | - |
| PCA yield [C–mmol C–mmol<sup>-1</sup>] | 0.02 | 0.04 | 0.06 | 0.33 | 0.10<sup>b</sup> | 0.19<sup>b</sup> |

<sup>a</sup>Following the additional D-glucose feed in this condition the resulting dynamics of measured process quantities were too complex for fitting of a valid process model.
<sup>b</sup>Calculated from the total amount of formed PCA divided by the total amount of consumed D-xylose.

**Fig. 2.**
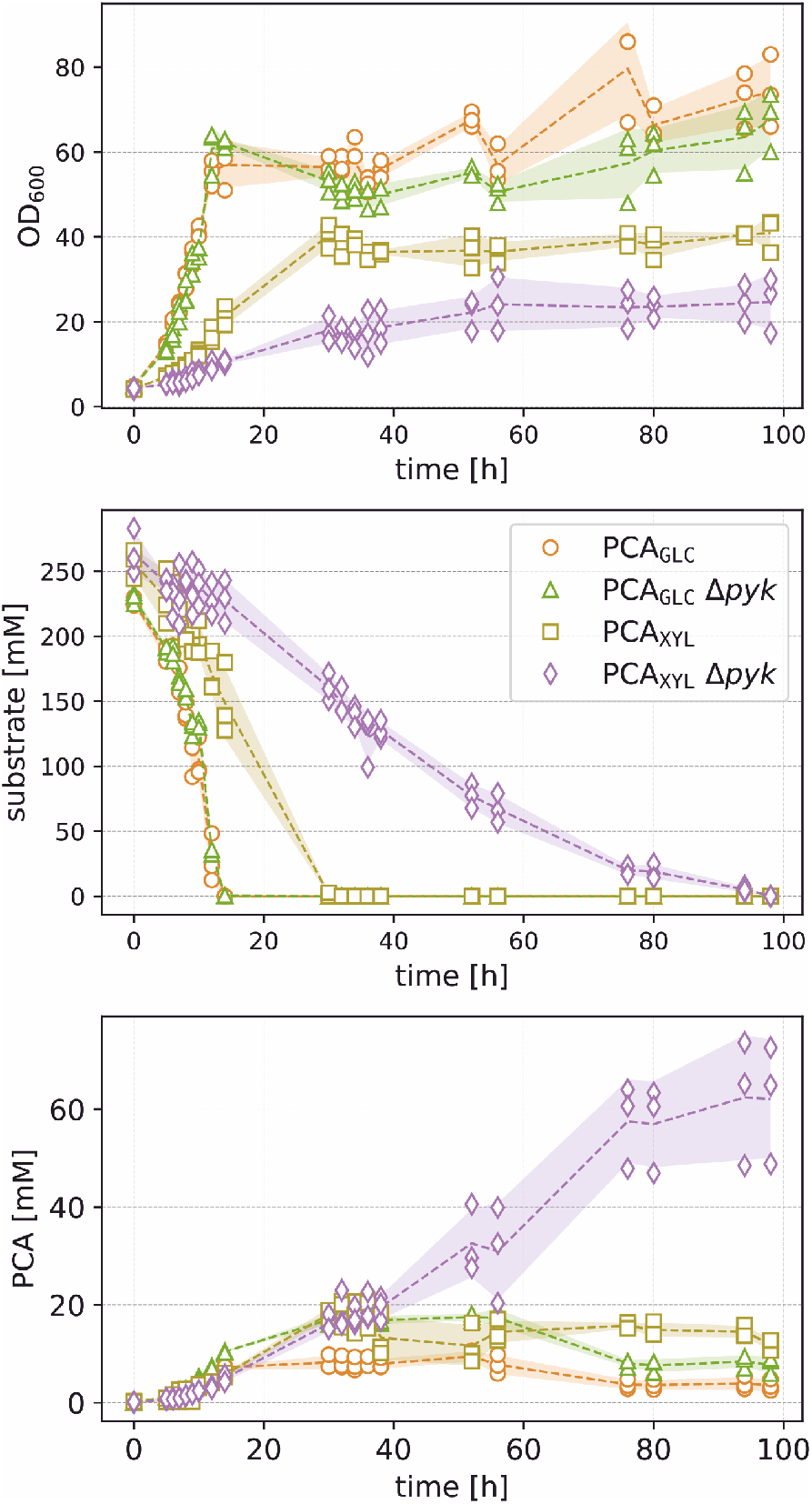
Comparative phenotyping of engineered *C. glutamicum* PCA producer strains. Shake flask cultivations were performed in defined CGXII medium, supplemented with either 222 mM d-glucose (PCA_GLC_, PCA_GLC_ Δ*pyk*) or 266 mM d-xylose (PCA_XYL_, PCA_XYL_ Δ*pyk*) as sole carbon and energy source. Mean values (dashed lines) and standard deviations (shaded areas) were derived from the three depicted independent cultures.

As expected, the inactivation of pyruvate kinase has no effect on growth of the d-glucose-based PCA producer strain. For *C. glutamicum* wild type and the corresponding *pyk*-deletion mutant it was already demonstrated that growth was not affected in defined CGXII medium with d-glucose as sole carbon and energy source [42]. The high demand of pyruvate and further pyruvate-derived precursors for biomass production might be completely fulfilled by the PTS-coupled d-glucose uptake. Alternatively, pyruvate could be formed via the concerted action of PEP carboxylase, malate dehydrogenase and malic enzyme, as shown previously for a pyruvate kinase-deficient wild-type strain of *E. coli* cultivated under d-glucose-limiting conditions [43].

By contrast, the additional inactivation of pyruvate kinase in the strain PCA_XYL_ resulted in a strong reduction of the specific growth rate (cf. Table 2). The observed slow, but steady growth of strain PCA_XYL_ Δ*pyk* up to an OD_600_ ≈ 25 during shake flask cultivation can only be explained by an alternative flux mode for pyruvate supply involving the operation of malic enzyme. In fact, a similar role of malic enzyme was shown for *C. glutamicum Δpyk* when grown on different gluconeogenetic substrates [44]. From our GC-ToF-MS analysis, we found extracellular accumulation of malate exclusively with strain PCA_XYL_ *Δpyk*, further supporting this hypothesis (see Supplementary Information).

The combined inactivation of pyruvate kinase and introduction of the non-PTS substrate d-xylose in strain PCA_XYL_ Δ*pyk* led to the observed superior PCA production performance. This is mainly due to the higher availability of both PCA precursors, E4P and PEP, which also resulted in a significantly higher accumulation of the direct condensation product DAHP compared to all other strains (see Supplementary Information).

### Development of a one-pot PCA production process

Process development for PCA production was initiated by cultivating strain PCA_XYL_ Δ*pyk* in a parallel bioreactor system (1.2 L) in defined CGXII medium containing 1 % w v^−1^ d-glucose as sole carbon and energy source for biomass growth, followed by repeated pulse-feeding of d-xylose to foster PCA formation (Figure 3).

**Fig. 3.**
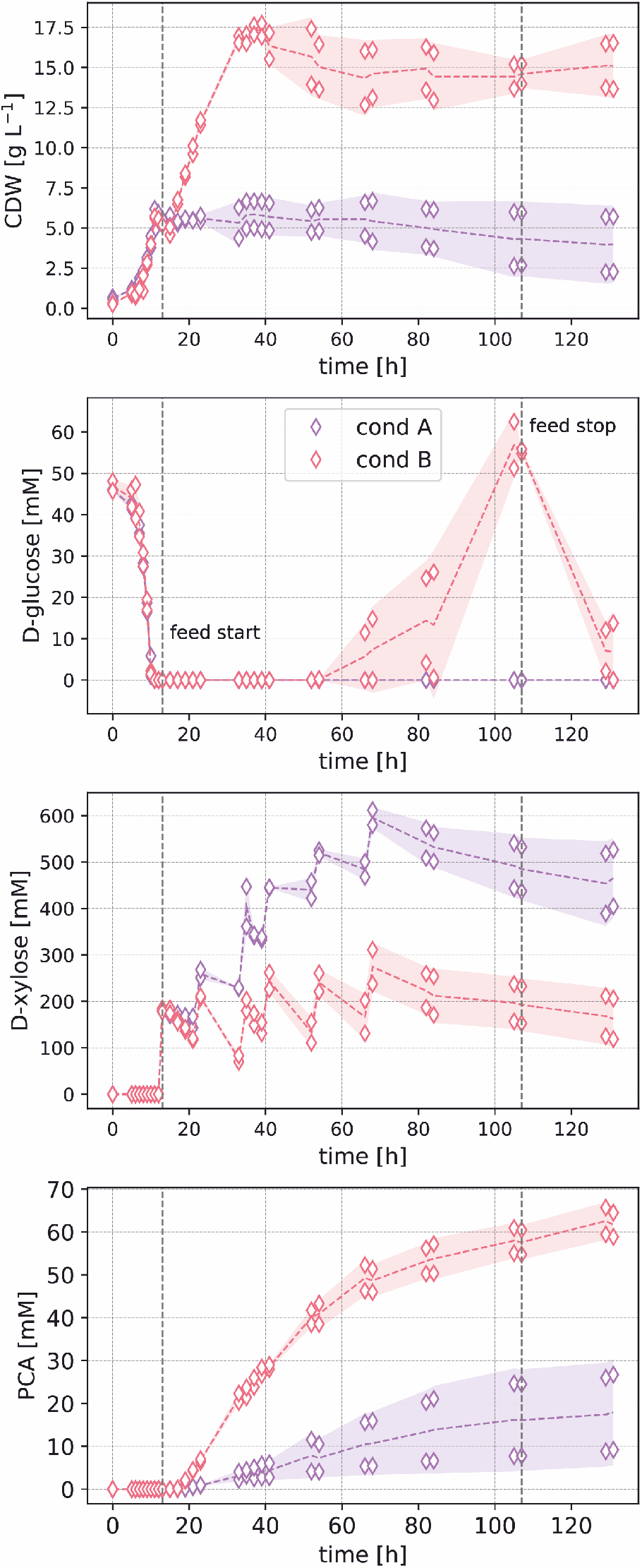
Fed-batch bioreactor cultivations of *C. glutamicum* PCA_XYL_ Δ*pyk* strain in CGXII medium containing 10 g L^−1^ d-glucose for initial batch growth. In condition A, starting at *t* = 13 h, a total of 162 g d-xylose was pulsed over the whole growth-decoupled production phase. In condition B an additional feed of 0.75 g h−1 d-glucose was introduced. d-Glucose feeding was stopped after 107 h of cultivation to avoid its further accumulation. Mean values (dashed lines) and standard deviations (shaded areas) were derived from the two depicted independent cultures.

Within the first 13 h of cultivation, d-glucose was completely consumed and the cell dry weight increased to 5.5 ± 0.1 g L^−1^. Subsequently, both plasmids (i.e. for d-xylose assimilation and PCA formation) were induced by IPTG and d-xylose feeding was initiated. As a result, the cells produced PCA from d-xylose in a growth-decoupled manner. During the cultivation, the d-xylose conversion and PCA formation rate significantly slowed down, resulting in titers of 9.2 mM and 26.7 mM in both replicates, respectively.

The decrease of the PCA formation rate towards the end of the cultivation could be attributed to the reduced activity of 3-dehydroshikimate dehydratase (QsuB). Previous *in vitro* analysis of the substrate-saturated QsuB enzyme from *C. glutamicum* showed non-competitive inhibition (**K*_I_* = 0.96 mM) at a concentration of 0.6 mM PCA [45]. Further *in vitro* analysis of this enzyme showed its highest activity in presence of 1 mM Co^2+^ and at a pH of 8.0 ± 8.4. Although both factors are considered not physiologically optimal for the cultivation of *C. glutamicum*, this bacterium is still able to grow at extracellular concentrations of 2 mM Co^2+^ [46] and establish pH homeostasis over a pH range from 6.5 ± 8.0 [47]. Therefore, strategies such as *in situ* product removal and media optimization could presumably maintain the enzymatic stability required for efficient PCA production over longer cultivation periods. However, no clear distinction could be made from our data regarding the physiological reasons for the decreased activity of 3-dehydroshikimate dehydratase. Therefore, the total loss of enzyme activity is modeled by assuming constant enzyme degradation and this approach resulted in a good description of the observed PCA dynamics (see Supplementary Information).

In order to achieve a higher PCA productivity with strain PCA_XYL_ Δ*pyk*, an additional d-glucose feed of 0.75 g_GLC_ h^−1^ after the initial batch phase was included. As expected, this resulted in an additional and substantial increase in biomass production up to 17.30 ± 0.41 g L^−1^ within 40 h of cultivation before some other essential media components became limiting for growth. In the subsequent growth-decoupled produc-tion phase, the final PCA titer was increased to 61.7 ± 4.0 mM (9.5 ± 0.6 g L^−1^). The observed higher d-xylose conversion rate to PCA was likely due to the increased biomass that remained metabolically active during assimilation of d-glucose. The microbial production of PCA from d-glucose has been previously reported in various bioreactor fermentations using engineered *C. glutamicum* strains. A production titer of 1.1 g L^−1^ PCA was reported after 96 h of fed-batch cultivation using a mini-jar fermenter containing a complex medium [17]. Most recently, by following a two-step fed-batch approach, a much higher titer of 82.7 g L^−1^ PCA was realized [21]. In this approach, the engineered strain was first grown to high cell density using complex medium, and then the biotransformation step was started from 10 % w v^−1^ of the harvested biocatalyst. The industrial relevance of this process is questionable due to the use of higher-cost complex media for biocatalyst preparation. Moreover, the high biomass-to-volume ratio represents potential challenges for the subsequent cell separation and PCA purification. In our study, a one-pot fermentation process based on a defined and cheap cultivation medium was developed for reaching 1.7 % w v^−1^ cell density from d-glucose and a production of 9.51 g L^−1^ PCA from d-xylose.

### Separation and purification of PCA

Following the final one-pot production process, PCA was separated from the cell-free supernatants of two independent cultures R1 and R2. The applied process concept for the purification of PCA was adapted from a very recent study [37] and consists of a concentration step, an electrochemical pH shift, and a cooling crystallization step (Figure 4A). The latter two unit operations were splitted to ensure sufficient conductivity during the pH shift [50]. The used solid-liquid equilibria of PCA in the fermentation medium for different pH values is shown in Figure 4B. The course of the solubility over the pH of recent experimental data for 30 °C fits well with the results from the fitted Henderson-Hasselbalch equation [48, 49]. Based on the measured solubility of 5.0 g L^−1^ at 5 °C and a pH of 2.99, the equation was then used to estimate the solubility of PCA over the pH at a temperature of 5 °C. Initially, the concentration of supernatants from the two replicates (R1 and R2) amounted 9.0 and 9.9 g L^−1^, respectively. After the evaporation step, the concentration in both samples was increased to 26.4 and 30.1 g L^−1^, and the pH was measured as 5.11 and 4.83, respectively. Next, the electrochemical shift was induced and in both cases, the pH decreased linearly, while the concentration of PCA remained almost constant (Figure 4C). At the end of the shift the concentration in R2 slightly decreased, which may indicate leakage through the electrode cage. Since the initial pH of sample R1 was higher than that of R2, the required electrical charge was enlarged. Though, the linear slope is similar due to the nearly equal concentrations. The final pH was measured as 3.80 and 3.58, respectively.

**Fig. 4.**
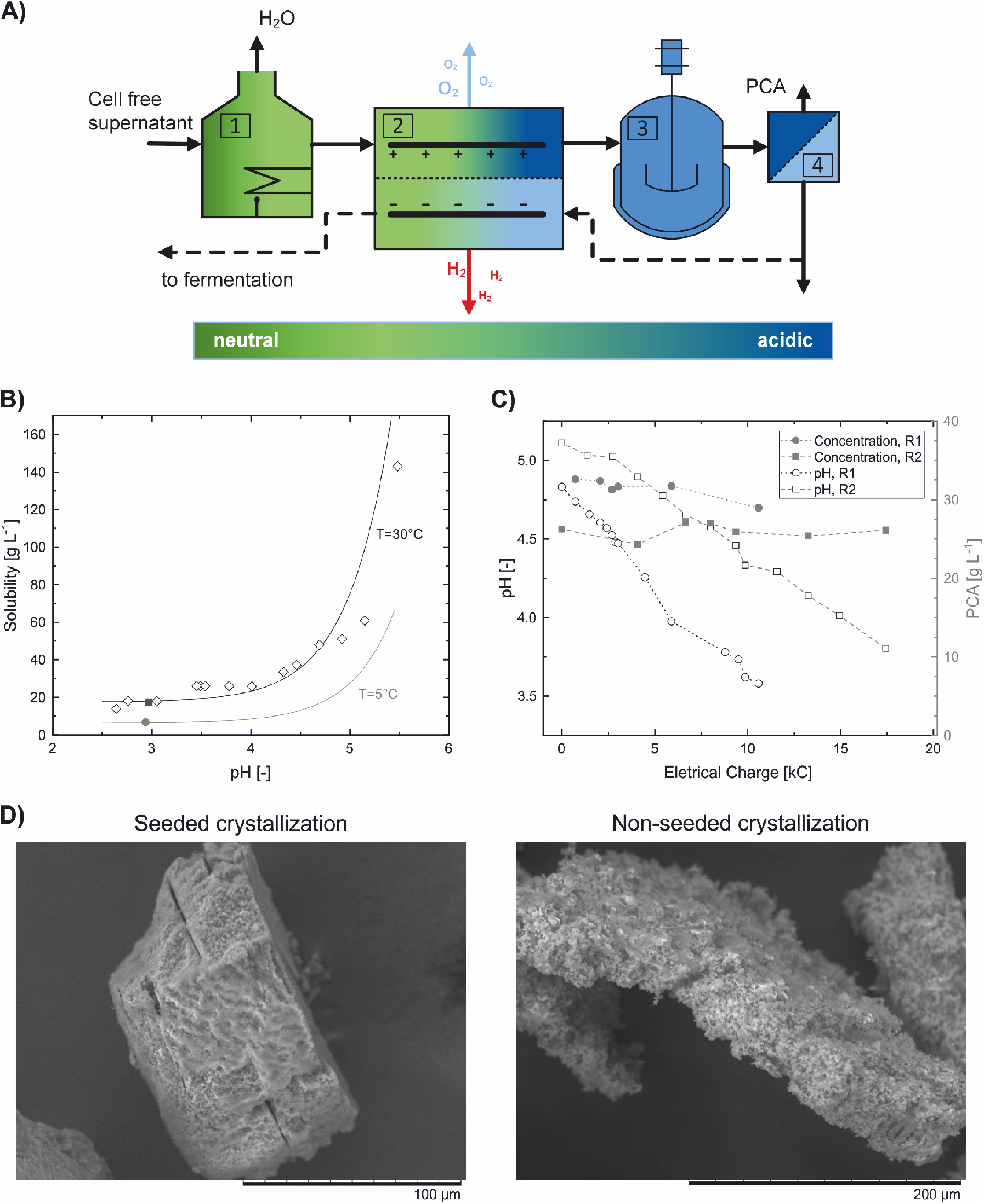
Separation of biotechnologically produced PCA. A) Process concept adapted from [37]. The supernatant is concentrated (1), the pH is electrochemical shifted (2), afterwards cooled to 5 °C (3) and filtered (4). Dashed line represents a possible recycle of the mother liquid to the fermentation. B) Solubility of PCA in fermentation medium from [48] (diamond) for 30 °C and experimental result for 5 °C (circle). The Henderson-Hasselbalch [49] equation was used to correlate pH-dependent solubility for 30 °C (black line) and 5 °C (gray line). C) Course of pH (black) and concentration of PCA (gray) during the electrochemical pH shift of fermentation medium A (circles) and B (squares). The average current in both experiments was *I* = 0.2 A. D) Scanning Electron Microscope pictures of the crystalline product of seeded (left) and non-seeded (right) crystallization captured with amplifications of 800 and 500, respectively.

The seeded and non-seeded crystallization experiments exhibited slow crystallization kinetics (Table 3). Even though the liquid was supersaturated roughly fivefold, the final concentration of the seeded crystallization still amounted 13.8 g L^−1^ after 14 days. Crystal nucleation in the non-seeded experiment was observed between seven and 14 days of experimental runtime. Such slow kinetics could be a consequence of the sample matrix, which may have contained sufficiently concentrated fermentation by-products to affect crystallization (see Supplementary Information).

**Table 3.** Results from the seeded and non-seeded crystallization experiments.

| Parameter/ condition | Seeded | Non-seeded |
| --- | --- | --- |
| Initial concentration [g L<sup>-1</sup>] | 27.4 | 27.4 |
| End concentration [g L<sup>-1</sup>] | 13.75 | 6.4 |
| Crystallization Yield [-] | 0.498 | 0.766 |
| Crystallization Efficiency $\xi$ [%] | 60.9 | 93.7 |
| Purity [wt %] | 95.4 | 91.8 |

The crystal surfaces of the seeded and non-seeded crystallization exhibited defects and showed the integration of small agglomerates (Figure 4D). This suggests that crystal growth was hindered and small PCA crystals formed agglomerates instead. Since the crystallization of PCA from the fermentation broth has not been studied before, this effect has to be investigated further. Sarma et al. (2014) detected different morphologies of PCA depending on the present temperature [29]. At temperatures below 10 °C, PCA forms needle-shaped crystals that have a tendency to break, agglomerate and de-celearate crystal growth [51]. Therefore, the crystallization of PCA at higher temperatures could be beneficial for the process.

Finally, the purity of the gained crystals was determined by HPLC for the seeded and non-seeded crystallization as 95.4 and 91.8 wt %, respectively.

## Conclusions

In this study, a highly sustainable bioprocess is presented for the microbial production and downstream processing of PCA. Combining *in silico* strain design with targeted metabolic engineering enabled the use of d-glucose and d-xylose as complementary carbon sources for cellular growth and product synthesis. The inactivation of pyruvate kinase and introduction of the non-PTS substrate d-xylose have significantly improved the PCA production performance in the engineered *C. glutamicum* strains. Purification of PCA was achieved by following a salt-free processing concept and yielded high-grade pure PCA crystals. With the established production and downstream processes, the sustainable biosynthesis of other hydroxybenzoic acids from alternative sugar feedstocks and their purification is within reach.

## Material and methods

### Model-based strain design and performance characterization

Flux balance analysis (FBA) was performed using a small-scale model of the central metabolism in *C. glutamicum*, which was additionally extended for the biosynthetic route of PCA (see Supplementary Information for more details). The modeling and visualization tool Omix was used for model definition and FBA was carried out using the available plug-in [52].

For the determination of key performance indicators (KPIs; including maximum titers, rates, and yields) of the different batch and fed-batch experiments conducted in this study, bioprocess modeling was performed using the pyFOOMB package [53]. The general bioprocess model and corresponding parameter estimates after fitting the model to the different experimental data sets can be found in the Supplementary Information.

### Bacterial strains, media, and growth conditions

All bacterial strains including their characteristics and sources are listed in Table 1. *E. coli* DH5*α* was used for cloning purposes and routinely cultivated aerobically at 37 °C either on Lysogeny Broth (LB) [54] agar plates (with 1.8 % w v^−1^ agar) or in reaction tubes filled with 5 mL LB medium on a rotatory shaker at 170 rpm. All *C. glutamicum* strains are derived from *C. glutamicum* ATCC 13032 [55] and were grown aerobically at 30 °C, either on Brain Heart Infusion (BHI) (Difco Laboratories, Detroit, USA) agar plates (with 1.8 % w v^−1^ agar) or in reaction tubes filled with 5 mL BHI medium on a rotatory shaker at 170 rpm. For strains harboring the cloning and construction vector pK19*mobsacB*, kanamycin was added to a final concentration of 25 μg mL^−1^ (*C. glutamicum*) or 50 μg mL^−1^ (*E. coli*). For the cultivation of *C. glutamicum* strains harboring the expression vectors pEKEx3 and pMKEx2, spectinomycin and kanamycin were supplemented to a final concentration of 100 μg mL^−1^ and 30 μg mL^−1^, respectively. Regulated gene expression was induced by the addition of isopropyl-*β*-d-thiogalactopyranoside (IPTG) to a final concentration of 1 mM.

### Plasmid and strain construction

All enzymes were purchased from Thermo Scientific (Schwerte, Germany). Standard protocols of molecular cloning, such as PCR and Gibson were used [56, 57]. *E. coli* DH5*α* was transformed via heat shock at 42 °C for 90 s with chemically competent cells prepared using the RbCl-method. *C. glutamicum* was transformed by electroporation followed by a heat shock at 46 °C for 6 min in BHIS medium (BHI medium supplemented with 90 g L^−1^ sorbitol). Regeneration of cells took place on a rotary shaker at 170 rpm (37 °C and 60 min for *E. coli*; 30 °C and 120 min for *C. glutamicum*) [58, 59]. The in-frame deletion of the *pyk* gene (cg2291) coding for pyruvate kinase in *C. glutamicum* was performed by two-step homologous recombination using the plasmid pK19*mobsacB-Δ pyk* according to a previously described protocol [60]. Verification of the constructed plasmid for gene deletion was performed by restriction analysis and deletion of *pyk* was verified by colony-PCR. For the used plasmids and oligonucleotides see Supplementary Information.

### Shake flask cultivations

Pre-cultures in 100 mL baffled shake flasks filled with 15 mL BHI were inoculated with single colonies from a fresh BHI agar plate and incubated for 8 h at 30 °C on a rotatory shaker at 250 rpm. These cultures were then used to inoculate a second pre-culture in 500 mL baffled shake flasks with 50 mL of defined CGXII medium [61] containing 4 % (222 mM) d-glucose as carbon and energy source. The incubation was performed for 15 h at 30 °C on a rotatory shaker at 250 rpm. Finally, the main culture was inoculated to an optical density at 600 nm (OD_600_) of 5.0 in 50 mL defined CGXII medium containing either 4 % (222 mM) d-glucose or 4 % (266 mM) d-xylose. Incubation was performed for 98 h at 30 °C on a rotatory shaker at 250 rpm. During cultivations, samples were taken for biomass and supernatant analysis at the indicated time points.

### Lab-scale bioreactor cultivations

Lab-scale cultivations were performed as biological duplicates using a parallel bioreactor system (Eppendorf/DASGIP, Jülich, Germany) with an initial working volume of 1.2 L. A pH of 7.0 was held constant during the cultivation by feeding 5 M H_3_PO_4_ and 5 M NH_4_OH on demand. Temperature and airflow were set to 30 °C and 0.5 vvm, respectively. Aerobic process conditions were maintained by controlling the stirrer speed (400 ± 1200 rpm) to achieve a dissolved oxygen concentration (DO) of at least 30 %. Online measurements were taken for pH (405-DPAS-SC-K80/225, Mettler Toledo), DO (VisifermDO 225, Hamilton) and exhaust gas composition (GA4, DASGIP AG). The cultivation started with 10 g L^−1^ d-glucose and inoculation was performed from an exponentially growing pre-culture using defined CGXII medium (40 g L^−1^ d-glucose) to an initial OD_600_ of 0.5.

After d-glucose was consumed (indicated by a sudden increase in the DO signal), IPTG was added to a final concentration of 1 mM and d-xylose pulsed-feeding was started (condition A). The d-xylose feed contained a solution of 450 g L^−1^ d-xylose in deionized water and a total feeding volume of 360 mL d-xylose solution was distributed into pulses to maintain the d-xylose concentration above 70 mM until the end of the cultivation. The d-xylose pulses of 71, 49, 60, 60, 60, and 60 mL were injected after 13, 23, 35, 41, 54, and 68 h of cultivation, respectively. For fed-batch fermentations, slow d-glucose feeding was started in addition to the d-xylose pulses (condition B). The d-glucose feed contained a solution of 500 g L^−1^ d-glucose in deionized water and the feed rate was set to 1.5 mL h^−1^. Feeding was performed for 94 h and was stopped after d-glucose started to accumulate.

### Biomass and supernatant analysis

Cell densities were assessed as OD_600_ measured using an UV-1800 spectrophotometer (Shimadzu, Duisburg, Germany). 1 mL cultivation broth was collected through a septum and diluted in 0.9 % w v^−1^ NaCl to an OD_600_ between 0.1 and 0.3. However, at 600 nm wavelength, PCA shows interference with the absorbance spectrum (see Supplementary Information). Therefore, all OD_600_ measurements were corrected by subtracting the OD_600_ value of the supernatant (obtained by sample centrifugation at 13 000 rpm for 10 min) from the OD_600_ value of the culture broth. For cell dry weight (CDW) determination 2 mL cultivation broth was collected in a weighted reaction tube, centrifuged at 13 000 rpm for 10 min and the resulting pellet was resuspended in 0.9 % w v^−1^ NaCl. After additional centrifugation, the supernatant was removed by decantation and the cell pellet was dried at 80 °C for 24 h followed by gravimetric CDW determination.

For substrate and product quantification, additional culture samples were centrifuged (13 000 rpm, 4 °C, 10 min) and the resulting supernatants were filtered through a cellulose-acetate syringe filter (0.2 μm, DIA-Nielsen, Düren, Germany). Separation of d-glucose, d-xylose, and PCA was performed on a HPLC system (Agilent 1100 Infinity, Agilent Technologies, Santa Clara, CA). The method used an Organic Acid Resin HPLC Column 300 × 8 mm (CS Chromatography, Düren, Germany) as stationary phase, 0.1 M H_2_SO_4_ with a flow rate of 0.6 mL min^−1^ as mobile phase, a column temperature of 55 °C and an injection volume of 20 μL. Detection of d-glucose and d-xylose was performed using a Refractive Index Detector at 35 °C. PCA was detected using UV light absorption at 215 nm with a Diode Array Detector. Concentrations were determined by applying an external standard in a weighted linear regression approach.

Untargeted metabolome analysis in culture supernatants was performed via an Agilent 6890N gas chromatograph coupled to a Waters Micromass GCT Premier high resolution time of flight mass spectrometer. Details regarding MS operation, sample preparation and peak identification are described in a previous study [62].

### Downstream processing of PCA

The pH-dependent solid-liquid equilibrium for PCA at 30 °C was taken from a previous study [48] and the solubility at 5 °C was determined using the shake-flask method [63]. The solubility of PCA was calculated for different pH values using solubility data, the *pK_a,COOH_* value of PCA, and the Henderson-Hasselbalch equation [49, 64].

Two samples from the replicate bioreactor cultivations applying condition B were centrifuged (8000 rpm, 45 min, 4 °C) and the supernatants were filtered (0.2 μm, Filtrox, Sankt Gallen, Switzerland) and concentrated (from 0.67 and 0.84 L to 0.25 and 0.28 L, respectively) at 101 °C at atmospheric pressure using a temperature-controlled magnetic hotplate stirrer with a ceramic plate (VWR, Langenfeld, Germany). The PCA concentration was measured using the HPLC analytics protocol described above.The pH was measured with a VWR pH-electrode pHenomenal® 110 (Langenfeld, Germany).

Afterwards, the pH of the concentrated samples was electrochemically shifted in a three-chamber electrolysis setup. In addition to the previously published setup [37, 65], the anode chamber was divided by inserting the anode into a cage consisting of polyethylene terephthalate glycol and a cation exchange membrane (Fumapem 14,100). This was done to avoid degradation of PCA at the anode, a phenomenon that was observed in preliminary experiments and previously published work [66]. A ruthenium oxide-coated titanium electrode (MAGNETO special Anodes B.V., Schiedam, Netherlands) and a nickel sheet (99 % purity) were used as anode and cathode, respectively. The temperature of all chambers was controlled at 30 °C and a magnetic stirrer was used for stirring the liquid in the anode chamber.

At the start of the experiment, the concentrated fermentation broth was filled into the anode chamber. 150 mL and 40 mL aqueous electrolyte, containing 0.5 M sodium dihydrogen phosphate dihydrate (NaH_2_PO_4_ · 2 H_2_O, purity *>* 98 %, Carl Roth, Germany), were added to the cathode chamber and the cage around the anode, respectively. During the experiment, the voltage was limited to 40 V and the current to 0.2 A. The pH was measured hourly and a sample was taken every two hours.

After the electrochemical pH shift, both liquids of the anode chamber were mixed and the pH and PCA concentration were measured. Finally, 100 mL medium were cooled to 5 °C and in seeded experiments, 0.32 g of sieved PCA seeds (*d*_50_ = 395 μm, *d*_10_ = 240 μm, *d*_90_ = 600 μm) were added to 350 mL medium and cooled to 5 °C. The crystallization efficiency was defined as the difference between start and end concentration divided by the maximal achievable crystal mass:

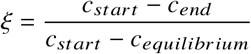

The identity and purity of the obtained crystals were analyzed using the described GC-ToF-MS and HPLC methods.

## Availability

The data used in this study can be made available upon reasonable request to the corresponding author.

## ACKNOWLEDGEMENTS

The authors acknowledge the financial support of the Bioeconomy Science Center as part of the projects HyImPAct (“Hybrid processes for Important Precursor and Active pharmaceutical ingredients”) and R2HPBio (“Renewables to high-performance bioplastics by sustainable production ways”). The scientific activities of the Bioeconomy Science Center were financially supported by the Ministry of Innovation, Science and Research within the framework of the NRW Strategieprojekt BioSC (no. 313/323-400-002 13).

## Supporting Information

### Model-based strain design and performance characterization

**Fig. S1.**
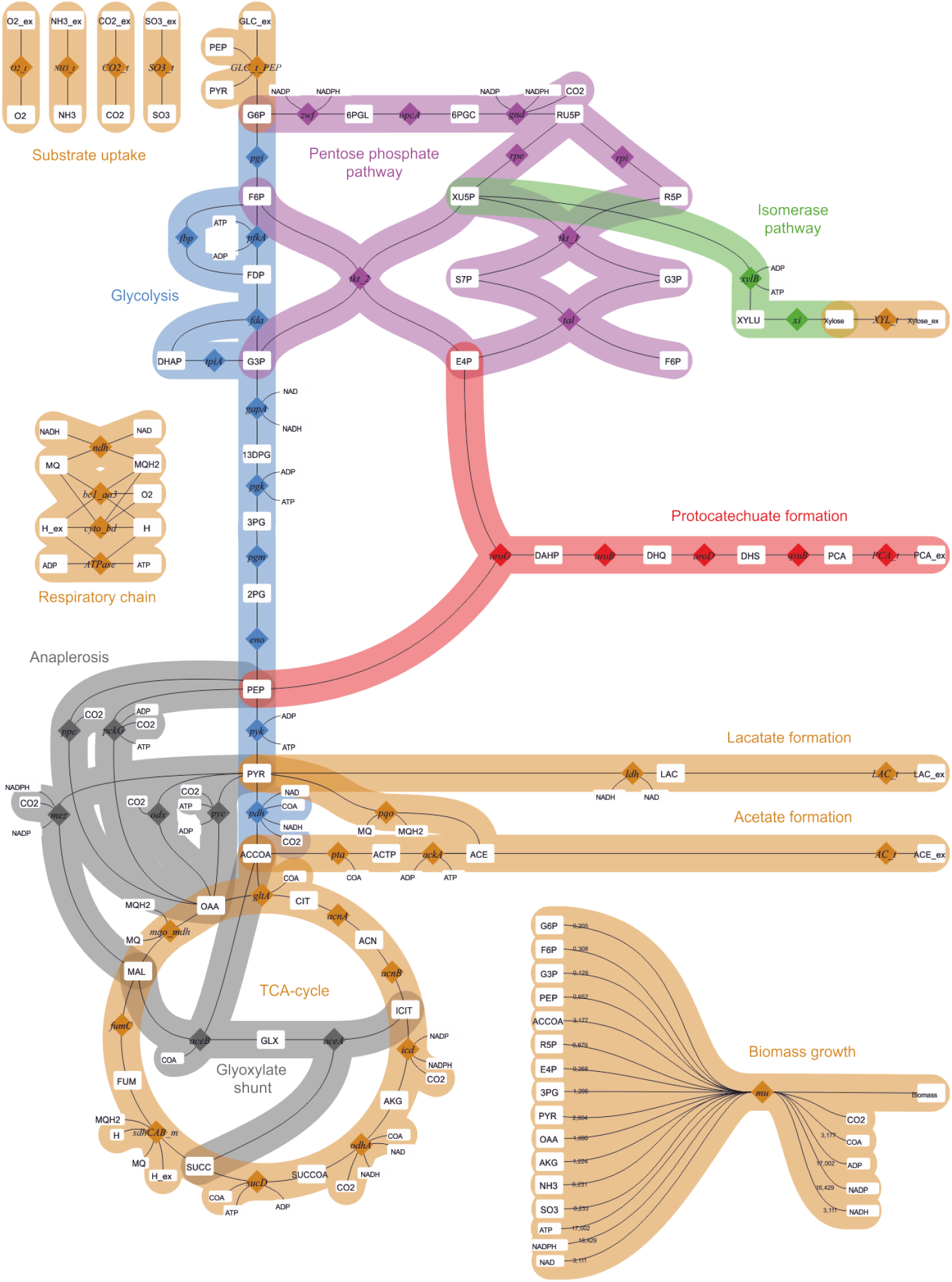
Metabolic network of *C. glutamicum* for *in silico* strain design. The isomerase pathway for d-xylose utilization was added to the set of native reactions of central metabolism.

For the determination of key performance indicators (KPIs) the following general bioprocess model was formulated:

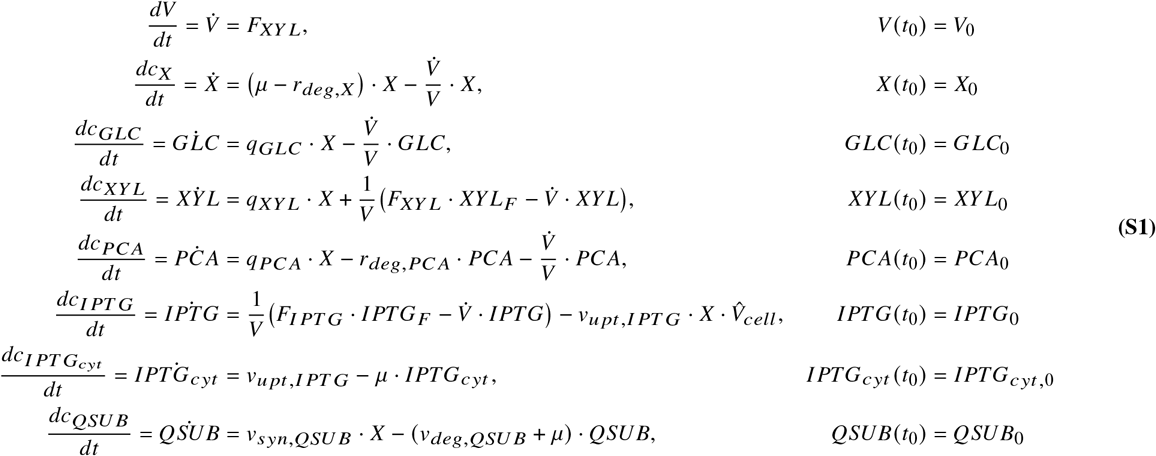

with

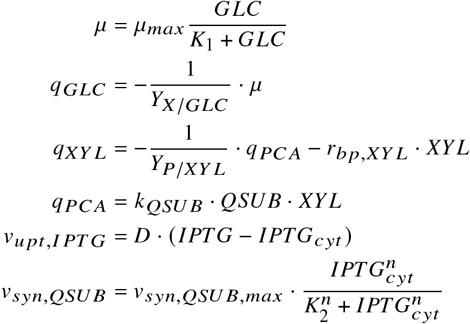

The concentration of biomass (*X*), d-glucose (*GLC*), d-xylose (*XYL*), protocatechuate (*PCA*) and extracellular inducer (*IPTG*) vary according to specific sources and sinks as well as in dependence of the changing working volume (*V*), following the feeding with d-xylose at the rate *F_XYL_*.

The growth rate of biomass (*μ*) is dependent on one limiting carbon source (here d-glucose) and modelled by classical Monod kinetics. The degradation rate of biomass (*r_deg,X_*) is modelled by mass action kinetics. The consumption rate of d-glucose (*q_GLC_*) is related to biomass growth by the yield coefficient *Y_X_|_GLC_*. The consumption rate of d-xylose (*q_XYL_*) is related to PCA formation by the yield coefficient *Y_P | XYL_* and additionally depends on the formation of byproducts at rate *r_*bp*,XYL_*.

The formation rate of PCA (*q_PCA_*) depends on one limiting carbon source (here d-xylose) as well as the activity of one limiting enzyme (here dehydroshikimate dehydratase (*QSUB*), encoded by the *qsub* gene. Plasmid-based expression of *QSUB* is induced by *IPTG* and modelled by hill kinetics. Only cytosolic *IPTG_cyt_* is active as inducer and the corresponding uptake into the cell with an average volume of *V_cell_* is modelled by diffusion kinetics. Finally, the PCA and enzyme degradation rates (*r_deg PCA_*, *v_deg, QSUB_*) are modelled by mass action kinetics.

Model formulation, parameter estimation and KPI determination was performed using the pyFOOMB tool [53]. The model in its full complexity was used to describe the dynamics of the fed-batch process with d-xylose feeding (Fig. S6). For the description of the other processes simplified versions of the model in Eq. S1 were derived that better suited the experimental conditions as well as given measurements (cmp. Figs. S2-S6).

Model parameters were estimated by running a repeated parameter estimation procedure, starting from different Monte-Carlo samples (so called “parametric bootstrapping”). In Table S1 the estimates for all model parameters across the different data sets are listed. Note, the integration of replicate experiments for parameter fitting is based on a differentiation between local (e.g., initial concentrations) and global (e.g., affinity constants) parameters (see [53] for detailed explanations).

**Fig. S2.**
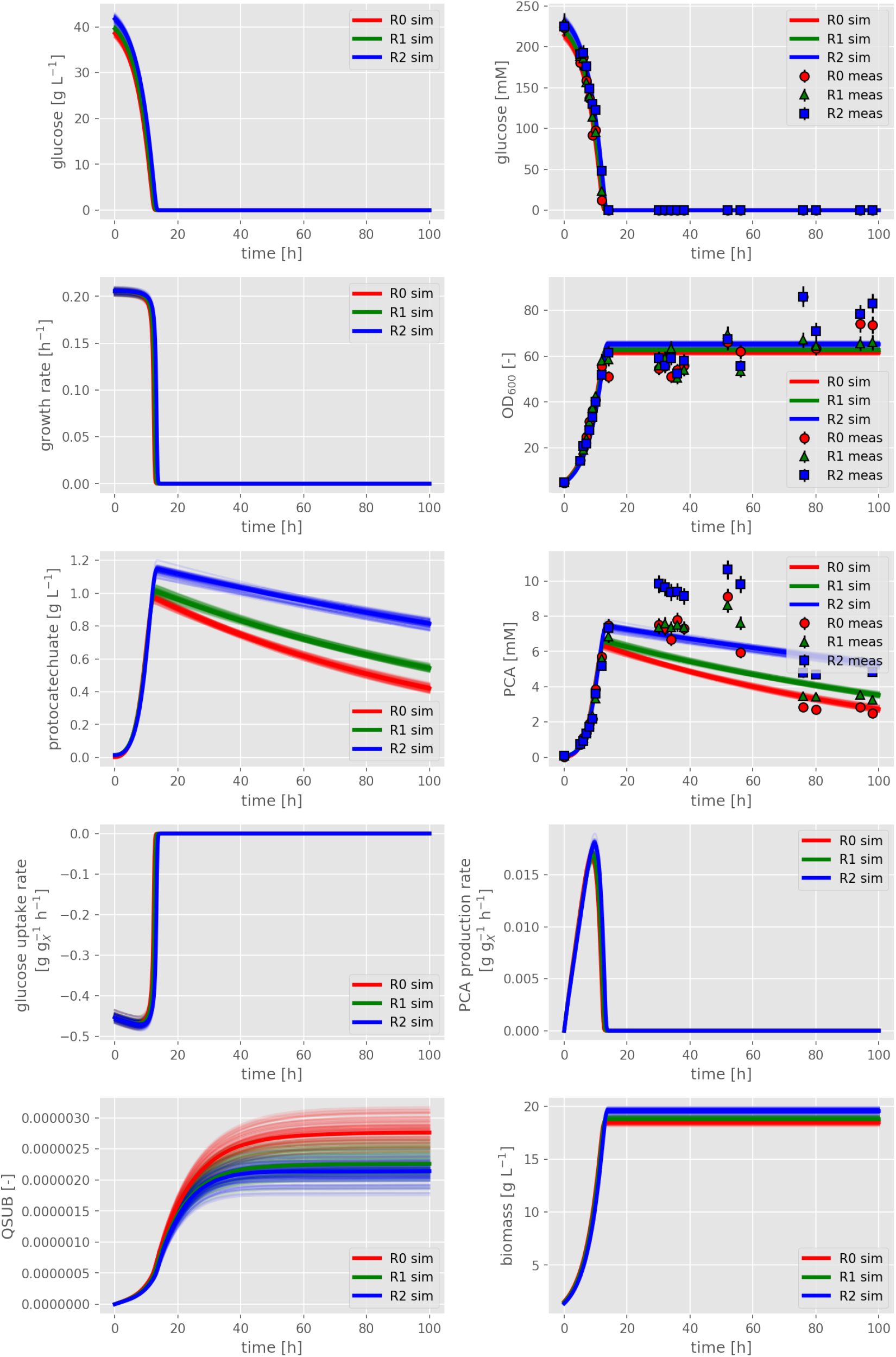
Data from replicate experiments (symbols) and corresponding model fits (straight lines) of strain PCA_GLC_ cultivated in shake flasks in defined CGXII medium, supplemented with 222 mM d-glucose as sole carbon and energy source.

**Fig. S3.**
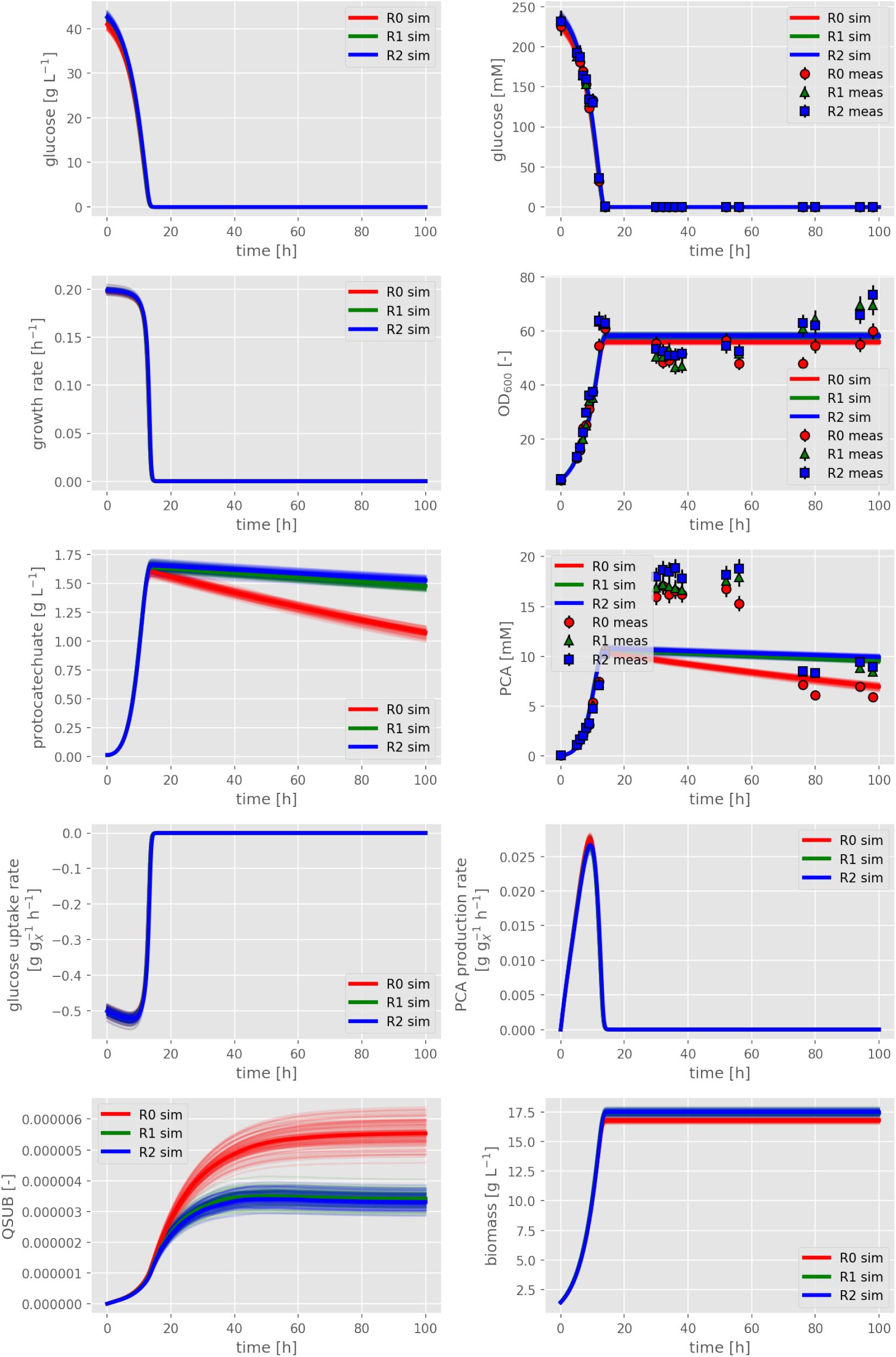
Data from replicate experiments (symbols) and corresponding model fits (straight lines) of strain PCA_GLC_ Δ*pyk* cultivated in shake flasks in defined CGXII medium, supplemented with 222 mM d-glucose as sole carbon and energy source.

**Fig. S4.**
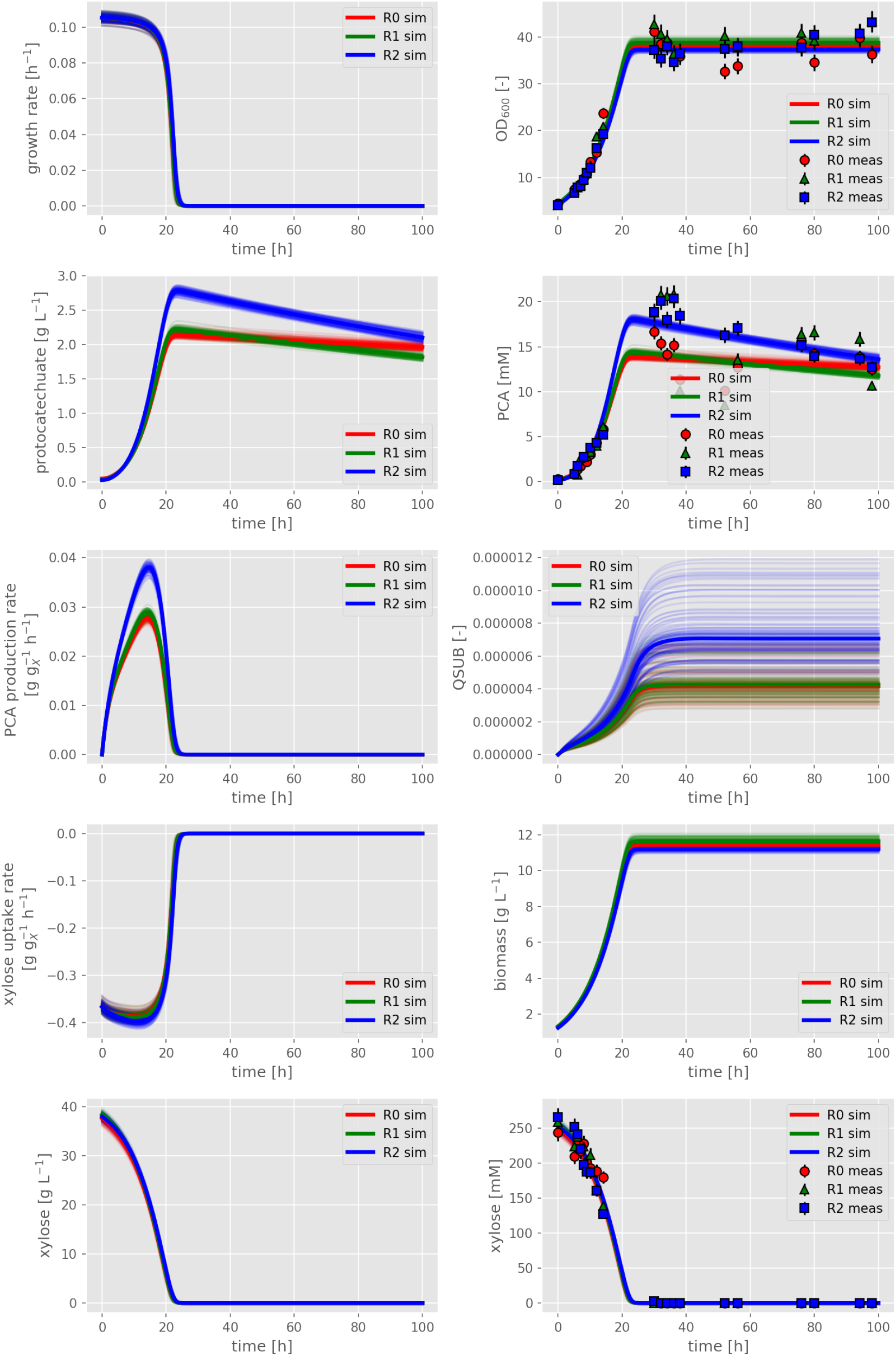
Data from replicate experiments (symbols) and corresponding model fits (straight lines) of strain PCA_XYL_ cultivated in shake flasks in defined CGXII medium, supplemented with 266 mM d-xylose as sole carbon and energy source.

**Fig. S5.**
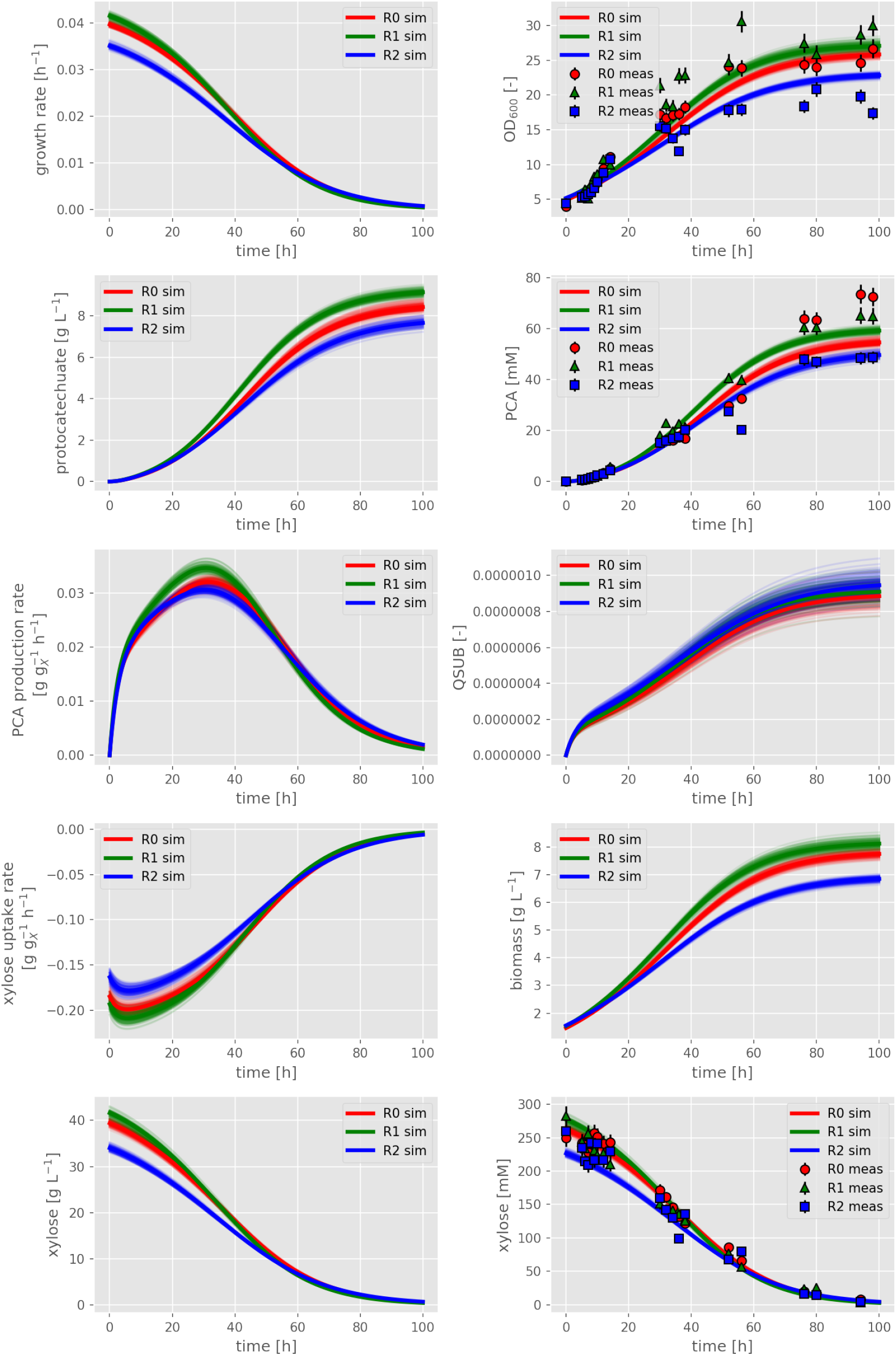
Data from replicate experiments (symbols) and corresponding model fits (straight lines) of strain PCA_XYL_ Δ*pyk* cultivated in shake flasks in defined CGXII medium, supplemented with 266 mM d-xylose as sole carbon and energy source.

**Fig. S6.**
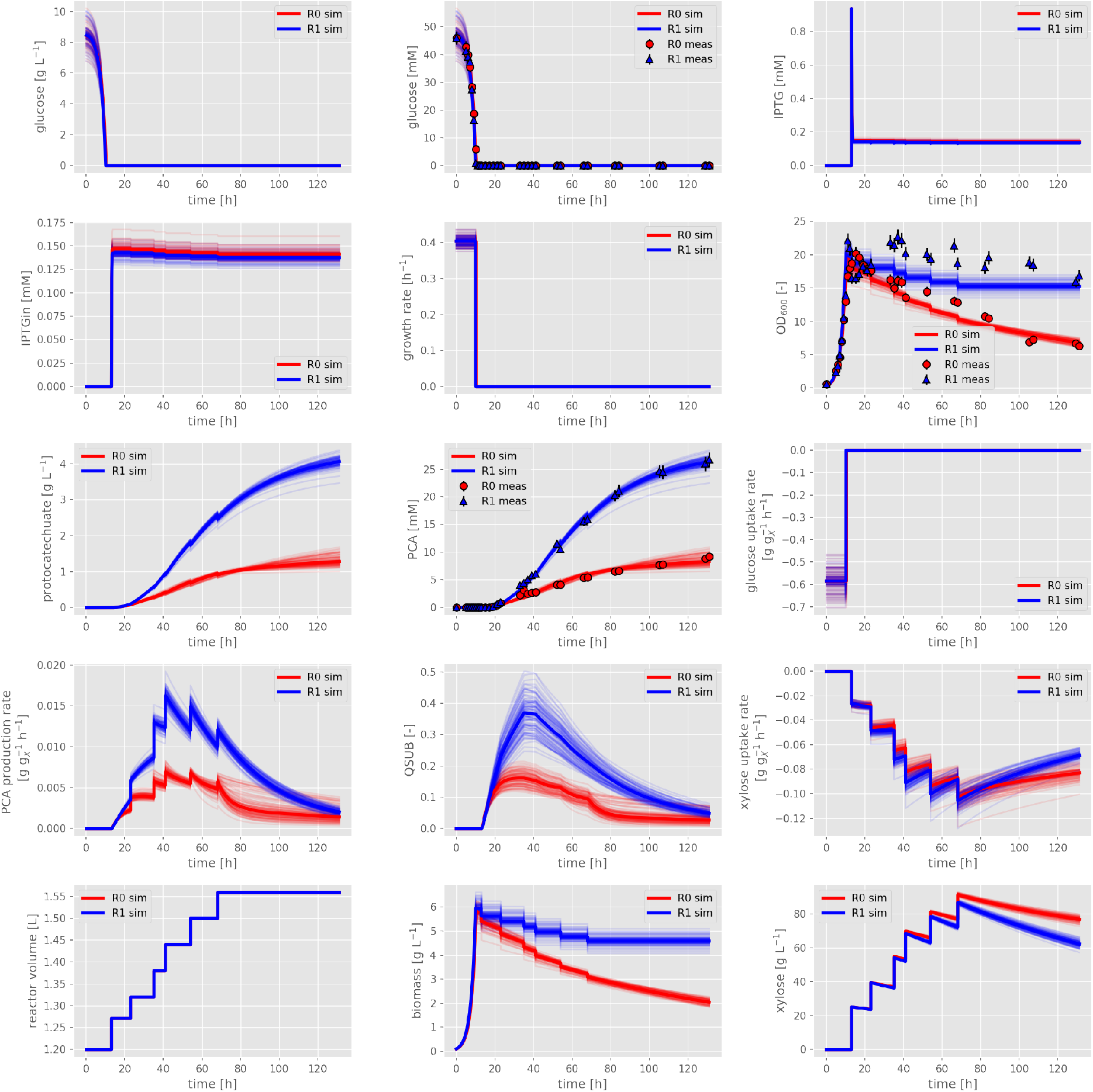
Data from replicate experiments (symbols) and corresponding model fits (straight lines) of strain PCA_XYL_ Δ*pyk* cultivated under fedbatch bioreactor conditions in defined GXII medium. For initial batch growth 10 g L^−1^ d-glucose was applied and at *t* = 13 h pulse feeding of d-xylose was started, providing a total amount of 162 g over the whole growth-decoupled production phase.

**Table S1.**
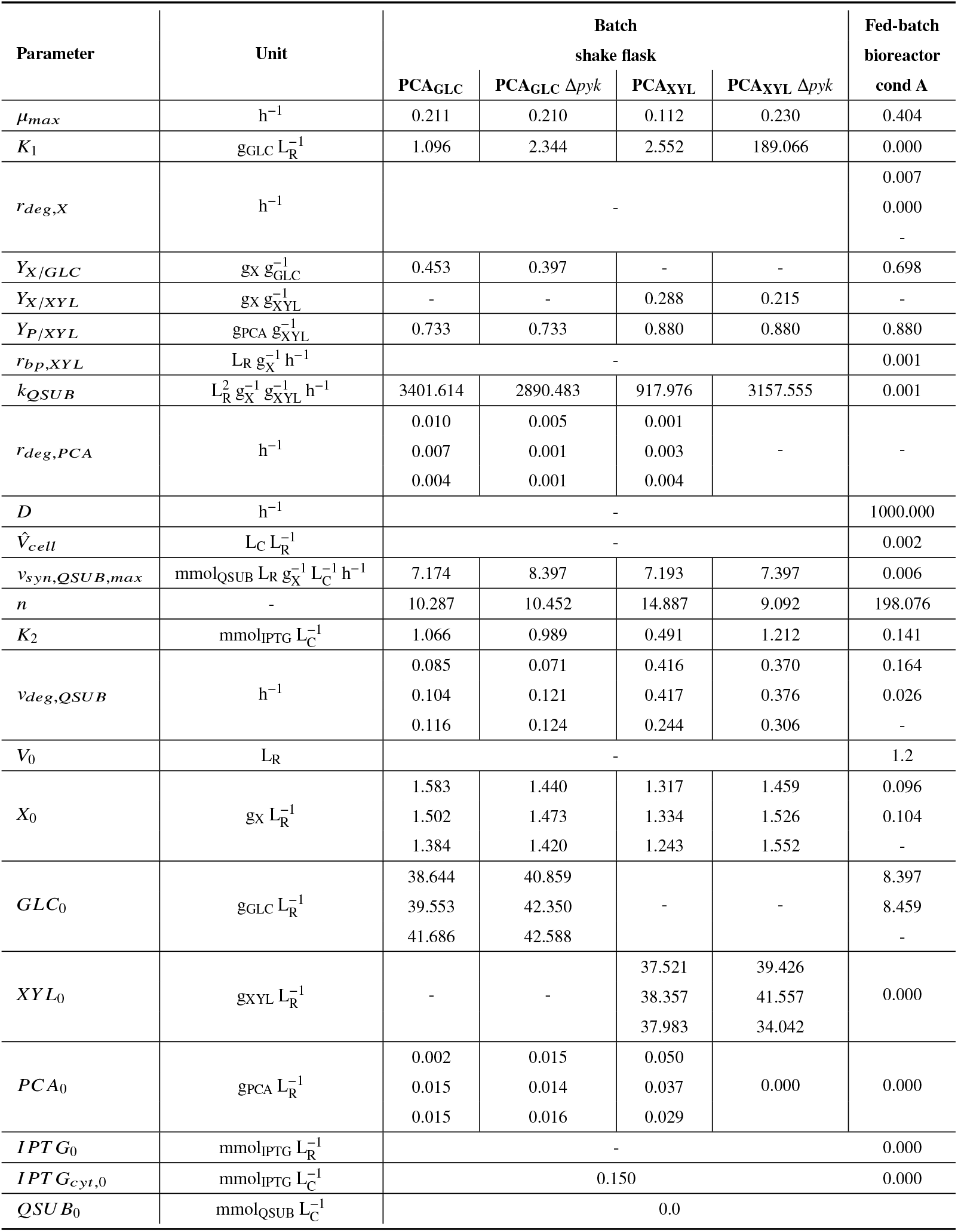
Parameter estimates of the bioprocess model of Eq. S1 based on the different cultivation experiments. Mean values from 100 runs of the parametric bootstrap are shown. Further statistical measures can be found in the available Jupyter notebooks.

| Parameter | Unit | Batch<br>shake flask |  |  |  | Fed-batch<br>bioreactor<br>cond A |
| --- | --- | --- | --- | --- | --- | --- |
| | | PCA<sub>GLC</sub> | PCA<sub>GLC</sub> $\Delta pyk$ | PCA<sub>XYL</sub> | PCA<sub>XYL</sub> $\Delta pyk$ | |
| $\mu_{max}$ | $h^{-1}$ | 0.211 | 0.210 | 0.112 | 0.230 | 0.404 |
| $K_1$ | $g_{GLC} L_R^{-1}$ | 1.096 | 2.344 | 2.552 | 189.066 | 0.000 |
| $r_{deg,X}$ | $h^{-1}$ | - | | | | 0.007<br>0.000<br>- |
| $Y_{X/GLC}$ | $g_X g_{GLC}^{-1}$ | 0.453 | 0.397 | - | - | 0.698 |
| $Y_{X/XYL}$ | $g_X g_{XYL}^{-1}$ | - | - | 0.288 | 0.215 | - |
| $Y_{P/XYL}$ | $g_{PCA} g_{XYL}^{-1}$ | 0.733 | 0.733 | 0.880 | 0.880 | 0.880 |
| $r_{bp,XYL}$ | $L_R g_X^{-1} h^{-1}$ | - | | | | 0.001 |
| $k_{QSUB}$ | $L_R^2 g_X^{-1} g_{XYL}^{-1} h^{-1}$ | 3401.614 | 2890.483 | 917.976 | 3157.555 | 0.001 |
| $r_{deg,PCA}$ | $h^{-1}$ | 0.010<br>0.007<br>0.004 | 0.005<br>0.001<br>0.001 | 0.001<br>0.003<br>0.004 | - | - |
| $D$ | $h^{-1}$ | - | | | | 1000.000 |
| $\hat{V}_{cell}$ | $L_C L_R^{-1}$ | - | | | | 0.002 |
| $v_{syn,QSUB,max}$ | $mmol_{QSUB} L_R g_X^{-1} L_C^{-1} h^{-1}$ | 7.174 | 8.397 | 7.193 | 7.397 | 0.006 |
| $n$ | - | 10.287 | 10.452 | 14.887 | 9.092 | 198.076 |
| $K_2$ | $mmol_{IPTG} L_C^{-1}$ | 1.066 | 0.989 | 0.491 | 1.212 | 0.141 |
| $v_{deg,QSUB}$ | $h^{-1}$ | 0.085<br>0.104<br>0.116 | 0.071<br>0.121<br>0.124 | 0.416<br>0.417<br>0.244 | 0.370<br>0.376<br>0.306 | 0.164<br>0.026<br>- |
| $V_0$ | $L_R$ | - | | | | 1.2 |
| $X_0$ | $g_X L_R^{-1}$ | 1.583<br>1.502<br>1.384 | 1.440<br>1.473<br>1.420 | 1.317<br>1.334<br>1.243 | 1.459<br>1.526<br>1.552 | 0.096<br>0.104<br>- |
| $GLC_0$ | $g_{GLC} L_R^{-1}$ | 38.644<br>39.553<br>41.686 | 40.859<br>42.350<br>42.588 | - | - | 8.397<br>8.459<br>- |
| $XYL_0$ | $g_{XYL} L_R^{-1}$ | - | - | 37.521<br>38.357<br>37.983 | 39.426<br>41.557<br>34.042 | 0.000 |
| $PCA_0$ | $g_{PCA} L_R^{-1}$ | 0.002<br>0.015<br>0.015 | 0.015<br>0.014<br>0.016 | 0.050<br>0.037<br>0.029 | 0.000 | 0.000 |
| $IPTG_0$ | $mmol_{IPTG} L_R^{-1}$ | - | | | | 0.000 |
| $IPTG_{cyt,0}$ | $mmol_{IPTG} L_C^{-1}$ | 0.150 | | | | 0.000 |
| $QSUB_0$ | $mmol_{QSUB} L_C^{-1}$ | 0.0 | | | | |

**Table S2.**
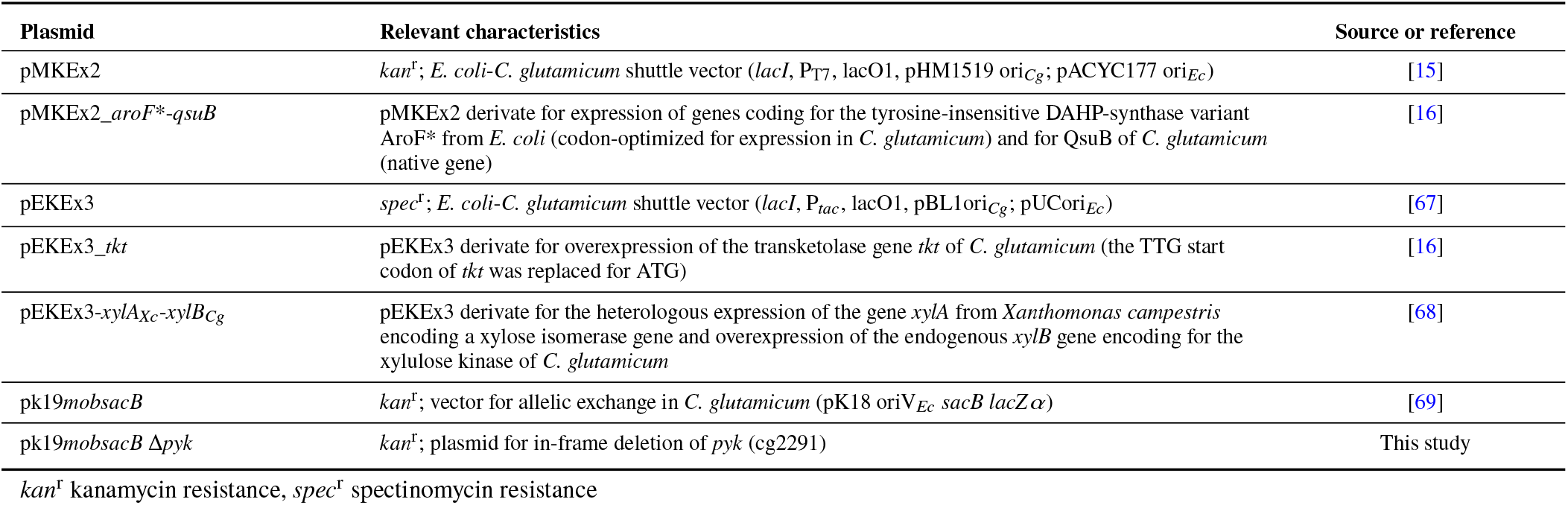
Plasmids used in this study.

**Table S3.**
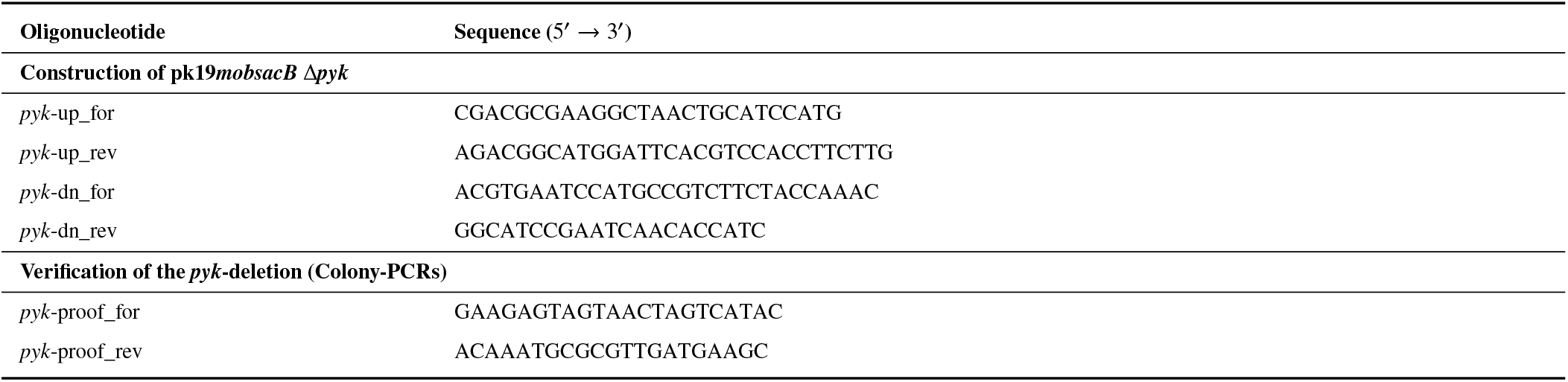
Oligonucleotides used in this study.

**Fig. S7.**
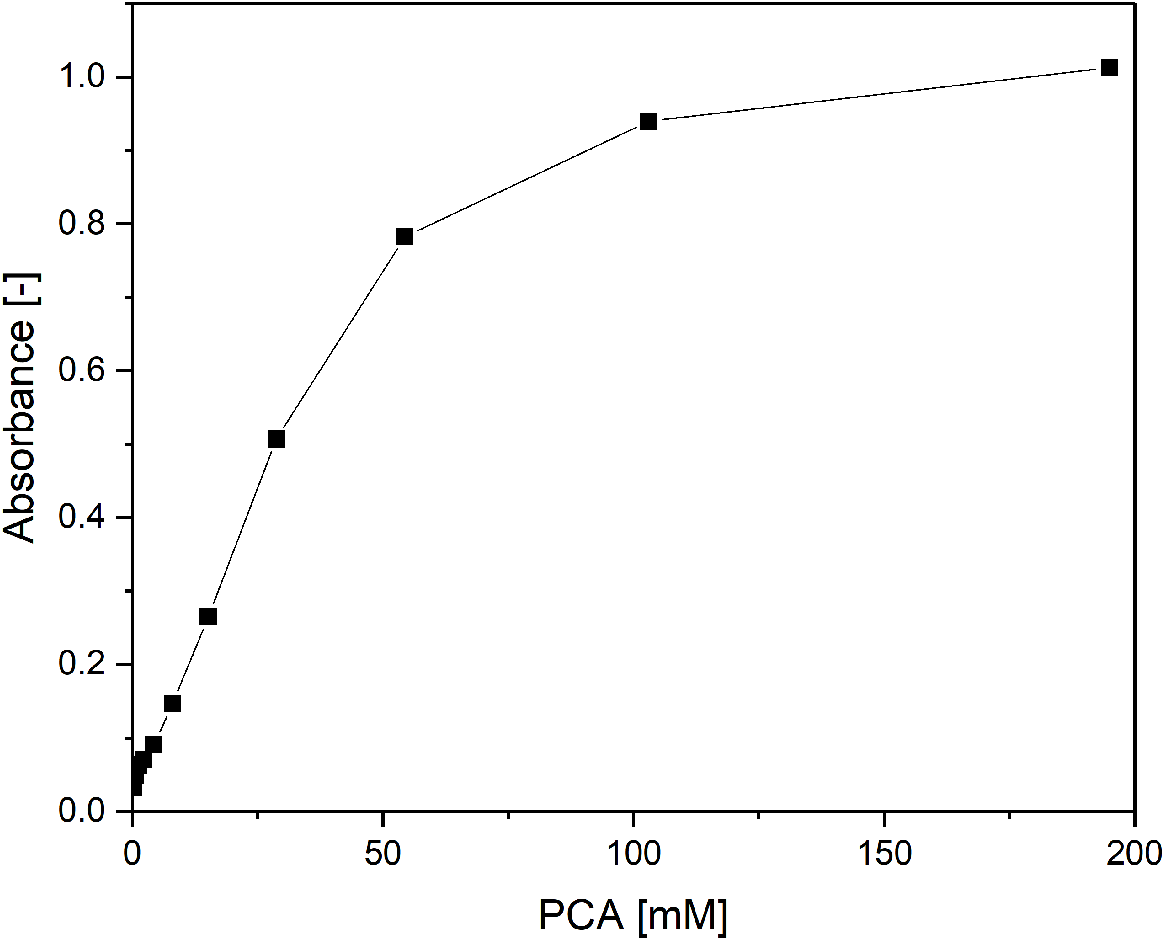
Absorbance of PCA at 600 nm and different concentrations in distilled water.

**Fig. S8.**
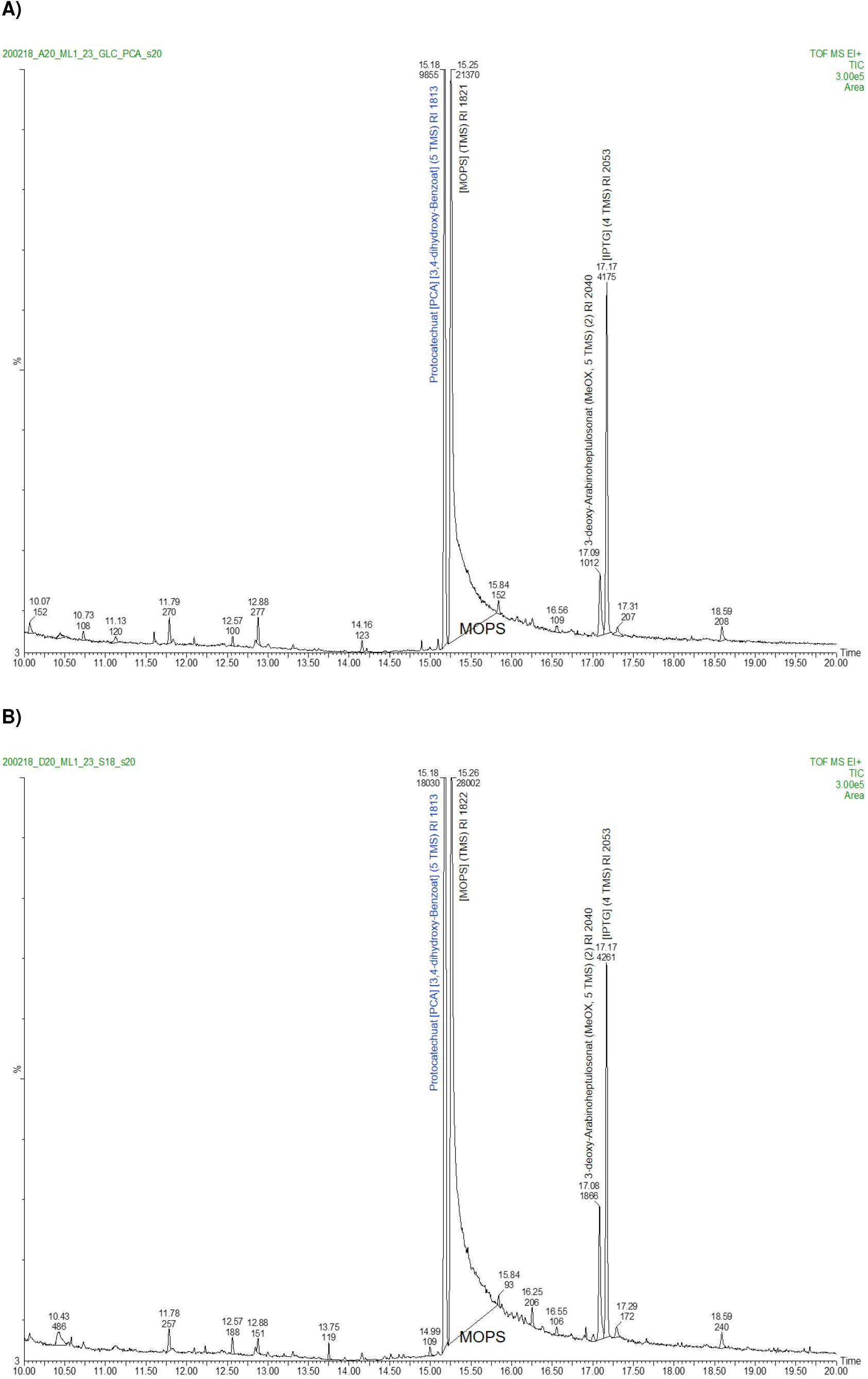
Selected chromatograms from untargeted metabolome analyis of culture supernatants from shake flask cultures of PCA producer strains: A) PCA_GLC_ and B) PCA_GLC_ Δ*pyk*.

**Fig. S9.**
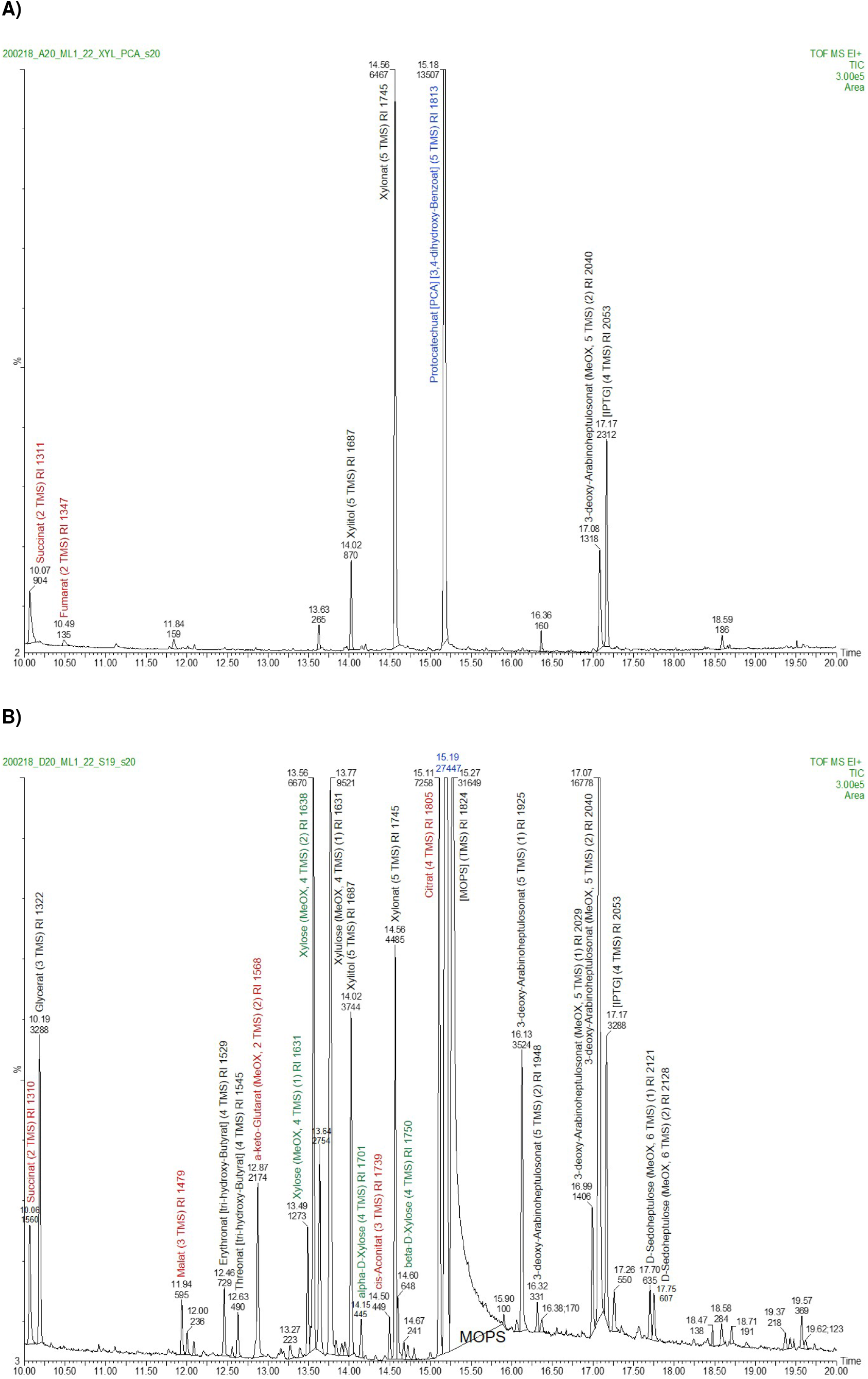
Selected chromatograms from untargeted metabolome analyis of culture supernatants from shake flask cultures of PCA producer strains: A) PCA_XYL_ and B) PCA_XYL_ Δ*pyk*.

